# Axonal mitochondria regulate gentle touch response through control of axonal actin dynamics

**DOI:** 10.1101/2024.08.13.607780

**Authors:** Sneha Hegde, Souvik Modi, Ennis W. Deihl, Oliver Vinzenz Glomb, Shaul Yogev, Frederic J. Hoerndli, Sandhya P. Koushika

**Affiliations:** Tata Institute of Fundamental Research, Homi Bhabha Road, Navy Nagar, Colaba, Mumbai-400005, India; Colorado State University, Anatomy and Zoology W309, 1617 Campus Delivery, Fort Collins, 80523 Colorado; Yale University, Boyer Center for Molecular Medicine, 295 Congress Ave, New Haven, CT 06510

**Author notes:** Institute of Clinical Anatomy and Cell Analysis, University of Tübingen, 72074 Tübingen, Germany (Oliver Vinzenz Glomb).

**Keywords:** Mitochondria, actin, *C. elegans*, plasma membrane proteins, RIC-7, SOD-2, UNC-9, MEC-4, MIRO, METAXIN

## Abstract

Actin in neuronal processes is both stable and dynamic. The origin & functional roles of the different pools of actin is not well understood. We find that mutants that lack mitochondria, *ric-7* and *mtx-2; miro-1*, in neuronal processes also lack dynamic actin. Mitochondria can regulate actin dynamics upto a distance ∼80 μm along the neuronal process. Absence of axonal mitochondria and dynamic actin does not markedly alter the Spectrin Membrane Periodic Skeleton (MPS) in touch receptor neurons (TRNs). Restoring mitochondria inTRNs cell autonomously restores dynamic actin in a *sod-2* dependent manner. We find that dynamic actin is necessary and sufficient for the localization of gap junction proteins in the TRNs and for the *C. elegans* gentle touch response. We identify an *in vivo* mechanism by which axonal mitochondria locally facilitate actin dynamics through reactive oxygen species that we show is necessary for electrical synapses & behaviour.

## Introduction

Actin in axons is both stable and dynamic in cultured hippocampal neurons and neurons *in vivo* (1-4). Stable actin consists of stationary actin-rich regions, the membrane periodic skelton (MPS) that consists of periodic actin rings and potentially other types of structures (1, 3, 4). The MPS comprises of Actin, Spectrin and Ankyrin arranged as regularly spaced rings ∼180 nm apart (3-5). The MPS provide mechanical support to axons, organizes membrane proteins e.g., sodium channels and regulates axonal microtubules (3, 6, 7).

The structure of dynamic actin in axons is unclear. The fomation of dynamic actin in hippocampal neurons in culture depends on Formin but is independent of the ARP 2/3 complex (1). The function of dynamic actin is critcal in neuron growth, branching and in axon regeneration, however its role in the adult axons is not clear. Dynamic actin in non-neuronal cells can be controlled by multiple pathways that likely converge on actin polymerizers like Formins and the ARP2/3 complex and depolymerzers like Gelsolin, Cofilin,etc (8-11). Signalling cascades activated by integrins, GPCRs, semaphorin 1A receptors, and organelles like mitchondria, endosomes regulate the actin cytoskeleton to drive process like cell shape changes, migration, proliferation (1, 8, 12-14). Studies from neurons in culture, neurons *in vivo* and non-neuronal cells suggest that mitochondria can regulate actin and acto-myosin structures during neuronal development and tissue remodeling (13-17). During development, mitochondria are implicated in synaptic elimination *in vivo* by promoting F-actin disassembly at synapses through the apoptotic pathway (15). By contrast, mitochondrially generated ATP supports F-actin patch formation in neurons in culture, helping neuronal branching and pre-synaptic assembly (16, 17). Additionally, tissue remodeling such as dorsal closure in *Drosophila* or wound healing of *C. elegans* epidermal cells, utilize the elevated mitochondrial calcium cell autonomously to remodel actomyosin structures (13, 14).

Prior work has shown a role for mitochondrial signaling and mitochondrial reactive oxygen species (ROS) signaling in regulating actin dynamics important for neuronal process development but its function in regulating actin dynamics in constitutive neuronal function is not well studied nor understood. In our study, we show that the axonal mitochondria regulate dynamic actin along the neuronal process via cytosolic ROS. Additionally using the TRN system we show that dynamic actin is important for distributions of several plama membrane proteins including gap junctions. The changes in distribuiton of these plasma membrane proteins might underlie the defective touch behavioural responses seen in mutants that lack dynamic actin.

## Results

### Mitochondria are present at actin rich regions in *C. elegans* touch receptor neurons

Mitochondria are present at actin enriched regions in many cell types including neurons (2, 17-20). To co-visualize F-actin and mitochondria in the sensory Posterior Lateral Mechanosensory (PLM neuron), we labelled mitochondria with mitochondrial matrix targeted GFP using mitochondrial localizing sequence-MLS (MLS::GFP) and actin with mCherry tagged to the calponin homology domain of the actin-binding protein utrophin (Utr-CH::mCherry) (1, 2).

As reported previously, we observe both stable actin that is stationary and dynamic actin in the neuronal processes of the PLM neuron (Fig. 1A, Movie 1). The stationary actin are present at a mean density= 52/100μm/min ±9.6 and polymerized actin patches apparent as trails in the kymograph have a mean density= 17/100μm/min ±4.5 (Fig. 1B). We observe stationary actin *in vivo* have varying lifespan and can be divided into a) short-lived actin (between 2 <120 secs) and b) long lived (> 120 secs) (Fig. 1C). In addition to trails that move in both anterograde and retrograde directions, dynamic actin consists of short-lived stationary actin hotspots and regions where actin shrinks (Fig. 1D, S1C). Most trails emerge from pre-existing stationary actin rich regions (∼ 53%) (Fig. 1E, F). Unlike in cultured hippocampal neurons, actin polymerisation did not show anterograde bias in the PLM neurons. The velocity, length and density of trails were similar in both anterograde and retrograde directions (Fig. S1C-H, Table S1). Compared to cultured neurons the actin trails had lower velocity and stationary actin and hotspots have shorter lifetimes (Fig. S1G, 1C) (1). Dual color time-lapse imaging of actin and mitochondria suggested that ∼90% of mitochondria are juxtaposed to actin-rich regions [Fig. 1 G,H (2)]. This suggests that mitochondria are present at actin rich regions along the neuronal process *in vivo*.

**Figure 1:**
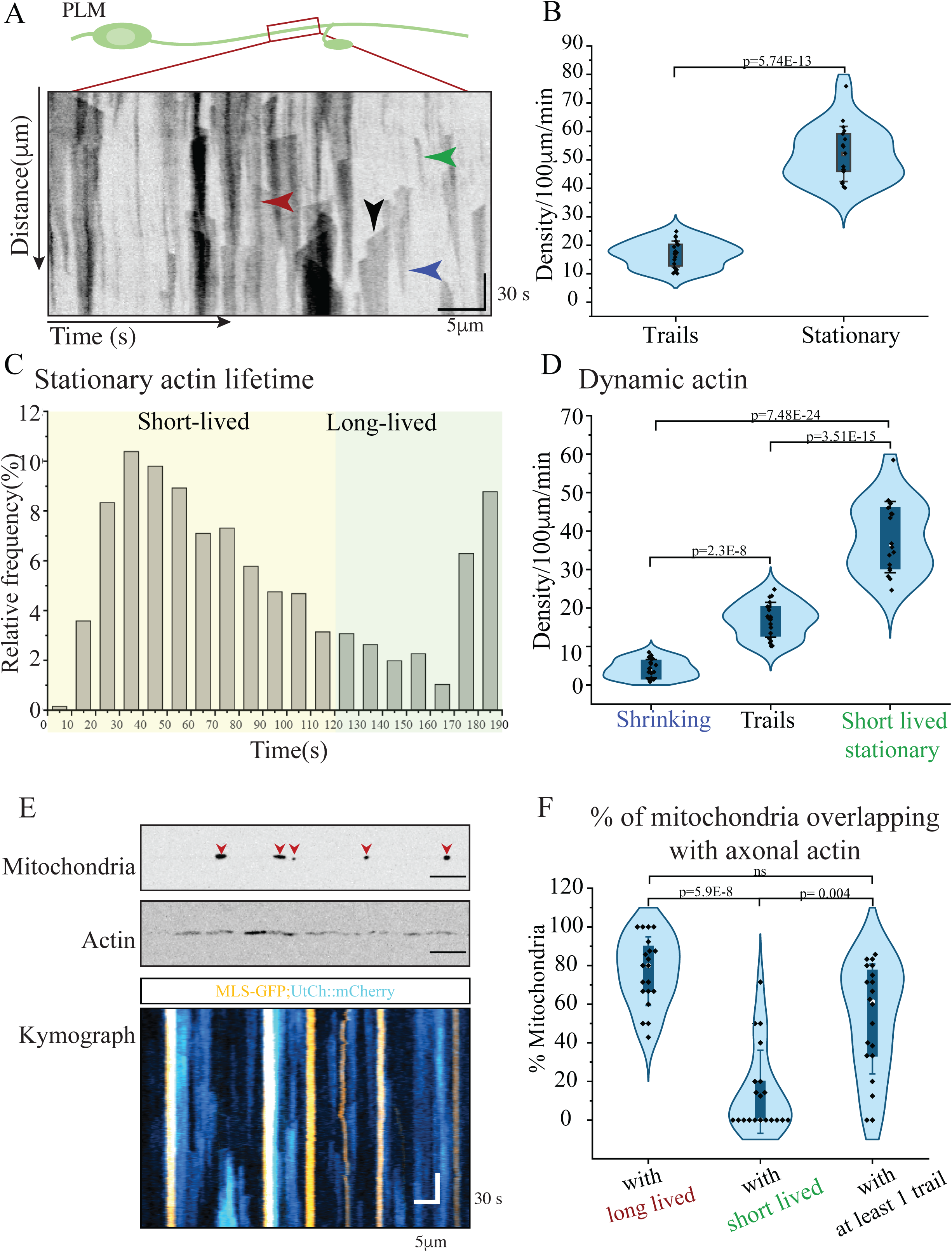
Mitochondria localize with actin in *C. elegans* TRNs. A) Kymograph obtained by time-lapse imaging of UtCH::GFP in PLM neuron. Arrow heads: black: trail, red: long-lived stationary actin, green: short-lived stationary actin, blue: shrinking actin region. Scale bar x axis=5μm, y axis=30 secs. B) Quantitation of the density of trails and stationary actin. n=20 animals for trails, 18 animals for stationary. Two-Sample t-test with unequal variance, Welch’s correction. C) Relative frequency of stationary actin over time. n=18 animals. D) Density of depolymerizing actin (shrinking), trails and stationary actin. n=20 animals for trails (510 events) and shrinking actin (130 events); n=18 animals for stationary actin (≥1300 events). One way ANOVA with Bonferroni correction. E) Representative images of mitochondria, actin and their corresponding kymographs obtained by dual color imaging. Scale bar x axis=5μm, y axis=30 secs. F) Quantitation of mitochondria juxtaposition with actin in the axons. n=20 animals. ns: non-significant. Kruskal Wallis ANOVA with Dunn’s Test

### Mitochondria are necessary and sufficient for dynamic actin along the neuron

Mitochondria and actin have been reported to influence each other (21-25). We thus investigated whether the presence of mitochondria affects actin in axons using *ric-7(lf)* animals and *metaxin; miro* double mutants where mitochondria are absent along neuronal processes of PLM but continue to be present in the cell body (Fig. S2A).

As expected, PLM major processes were devoid of mitochondria in *ric-7(lf)* and double mutants of *mtx-2(lf); miro-1(lf)* (Fig. 2B, C, D, H, S2A). We observe a strong reduction in the dynamic actin in the neuronal processes of PLM of *ric-7(lf)* and *mtx-2(lf); miro-1(lf)* as compared to wild type PLM neuronal processes (Fig. 2C, D, G and S2B, Movie 2, Movie 3, Tabls S2) and in the axons of HSN of *ric-7(lf))* (Fig., S2 D, E, Table S2). Long-lived stable actin was unaffected in these genotypes (Fig S2C, F), suggesting that mitochondria along the neuronal process are necessary for dynamic actin in the neuron. To determine if these effects were specific to actin and did not affect other cytoskeleton. We investigated microtubule dynamics by performing time-lapse imaging of EBP-2::GFP in *ric-7(lf)* (Fig. S2 G, H, E Movie 14). We observe similar fractions of plus end and minus end out comets (with respect to the cell body) in wild type and *ric-7(lf)* (Fig. S2 H, I).

**Figure 2:**
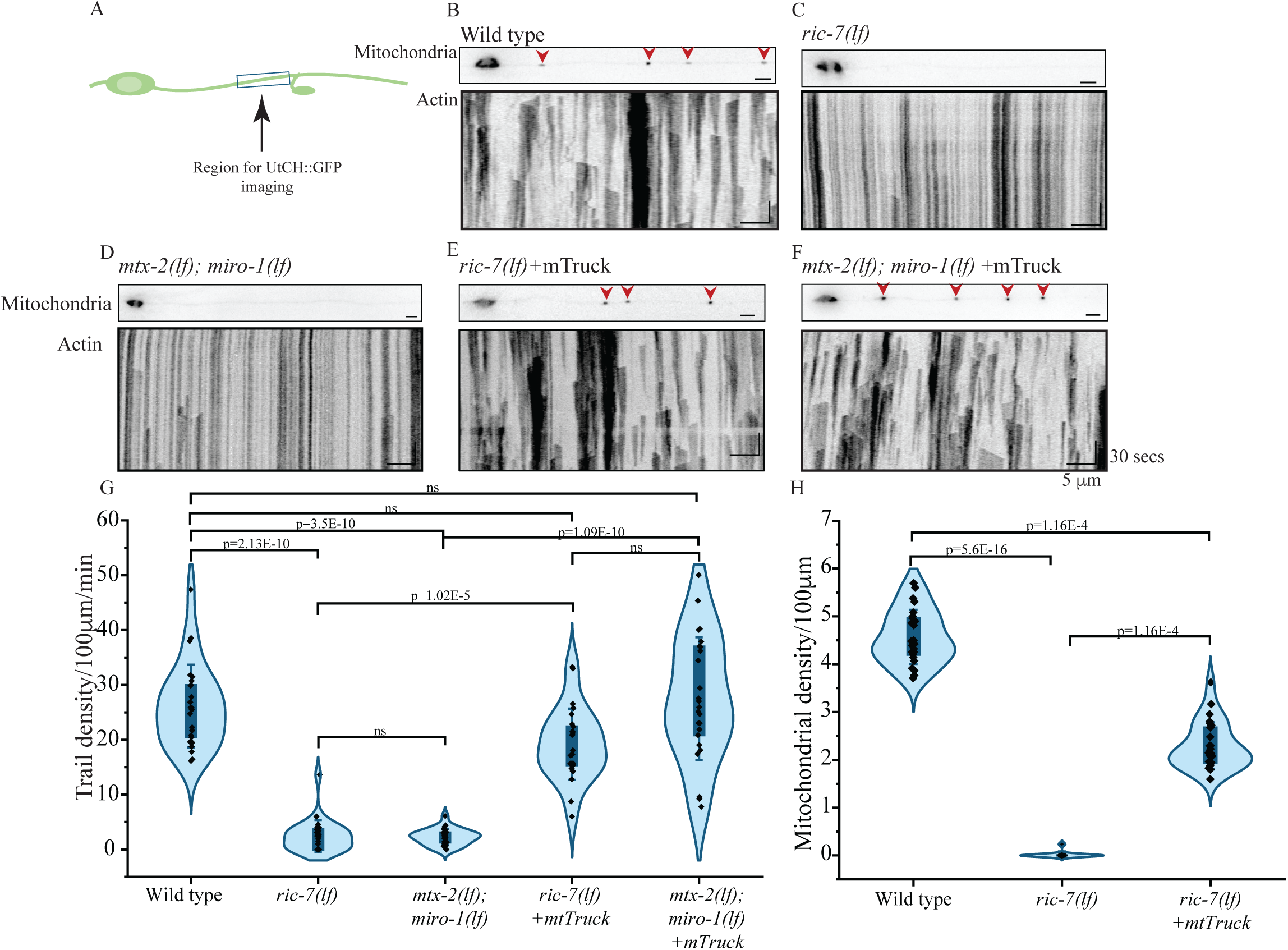
Mitochondria are necessary and sufficient for dynamic actin along the neuron. A) Schematic representation of the PLM neuron highlighting the region imaged for UtCH::GFP. B) Top panel: representative images of mitochondria in the neuron, bottom panel: representative kymographs for actin obtained by time lapse imaging of GFP::UtCH in wild type. Scale bar: x axis=5μm, y axis=30 sec C) Top panel: representative images of mitochondria in the neuron, bottom panel: representative kymographs for actin obtained by time lapse imaging of GFP::UtCH in *ric-7(nu447).* Scale bar: x axis=5μm, y axis=30 sec D) Top panel: representative images of mitochondria in the neuron, bottom panel: representative kymographs for actin obtained by time lapse imaging of GFP::UtCH in *mtx-2(gk444); miro-1(tm1966).* Scale bar: x axis=5μm, y axis=30 sec E) Top panel: representative images of mitochondria in the neuron, bottom panel: representative kymographs for actin obtained by time lapse imaging of GFP::UtCH in *ric-7(nu447) +tbEx307(*mTruck). Scale bar: x axis=5μm, y axis=30 sec F) Top panel: representative images of mitochondria in the neuron, bottom panel: representative kymographs for actin obtained by time lapse imaging of GFP::UtCH in *mtx-2(gk444);miro-1(tm1966)+tbEx307(*mTruck). Scale bar: x axis=5μm, y axis=30 sec G) Quantitation of trails density/100μm/min in wild type, *ric-7(nu447)*, *mtx-2(gk444); miro-1(tm1966)*, *ric-7(n447) +tbEx307(*mTruck), *mtx-2(gk444); miro-1(tm1966)+ tbEx307(*mTruck). n ≥25 animals for all genotypes, ≥ 50 trails for *ric-7(nu447)* and *mtx-2(gk444); miro-1(tm1966)*; ≥ 900 trails for other genotypes. Kruskal Wallis ANOVA with Dunn’s test. H) Quantitation of mitochondrial density/100μm in major process of PLM in wild type, *ric-7(nu447)* and *ric-7(nu447) +tbEx307(*mTruck), n≥ 25 animals. Kruskal Wallis ANOVA with Dunn’s test.

To restore mitochondrial distribution in the neuron independent of mitochondrial transport adaptors, we expressed Kinesin1 fused to mitochondrial protein TOM7 (mTruck) only in the touch receptor neurons (TRNs) (26). We observe that mitochondria are restored along the neuronal process in these animals but with a slightly reduced density (Fig 2E, F, S2A). In these animals, actin dynamics are restored similar to that seen in PLM in wild type animals (Fig. 2E, F, G, S2B) suggesting a cell-autonomous effect of axonal mitochondria on the presence of dynamic actin.

These results suggests that axonal mitochondria are necessary and sufficient for dynamic actin along the neuronal process.

### Mitochondria regulates actin dynamic regions locally

We investigated whether axonal mitochondria locally promote dynamic actin in the neuronal process or whether mitochondria can influence actin dynamics at a distance. We examined the extent of dynamic actin along the neuronal process in *ric-7* mutants where all mitochondria are restricted to the cell body (Fig 3A) and in *ric-7(lf)* expressing a mTruck transgene that does not restore mitochondria throughout the PLM neuronal process (Fig 3C).

**Figure 3:**
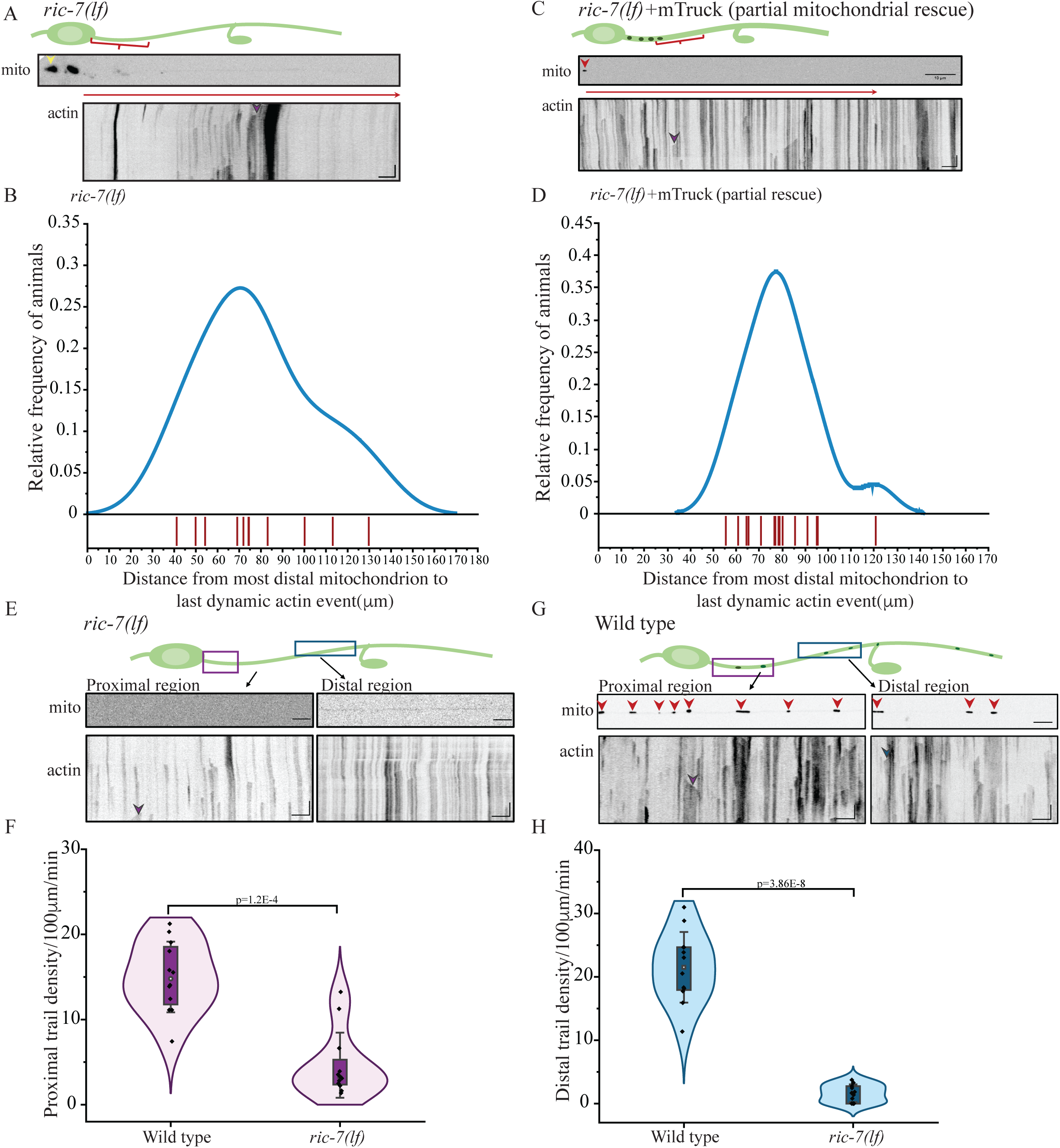
Mitochondria regulates actin dynamic regions locally. A) Top panel: representative images of mitochondria, bottom panel: representative kymographs for actin obtained by time-lapse imaging of GFP::UtCH at the proximal region of PLM major process in *ric-7(nu447).* Purple arrow head-representative trail. Scale bar: x axis=5μm, y axis=30 sec B) Quantitation of the distance of last dynamic actin seen in each animal (red vertical line) from the cell body in *ric-7(nu447).* Data plotted as rug plot. n>10 animals. C) Top panel: representative images of mitochondria, bottom panel: representative kymographs for actin obtained by time-lapse imaging of GFP::UtCH for the proximal region of PLM major process in *ric-7(nu447)+ tbEx371(*mTruck: incomplete mitochondrial rescue). Purple arrow head-representative trail. Scale bar: x axis=5μm, y axis=30 sec D) Quantitation of the distance of last dynamic actin seen in each animal (red vertical line) from the last mitochondrion in *ric-7(nu447) +tbEx371(*mTruck: incomplete mitochondrial rescue). Data plotted as rug plot. n>10 animals. E) Top panel: representative images of mitochondria, bottom panel: representative kymographs for actin obtained by time-lapse imaging of GFP::UtCH for the proximal and distal region of PLM major process of *ric-7(nu447).* Purple and blue arrow head-trail in proximal and distal region respectively. Scale bar: x axis=5μm, y axis=30 sec. F) Quantitation of trail density/100μm/min in proximal of *ric-7(nu447)* and wild type. n>10 animals for both genotypes. Mann-Whitney test. G) Top panel: representative images of mitochondria, bottom panel: representative kymographs for actin obtained by time-lapse imaging of GFP::UtCH for the proximal and distal region of PLM major process of wild type. Purple and blue arrow head-trail in proximal and distal region respectively. Scale bar: x axis=5μm, y axis=30 sec H) Quantitation of trail density/100μm/min in distal region of *ric-7(nu447)* and wild type. n>10 animals for both genotypes. Two-Sample t-test with unequal variance, Welch correction.

In *ric-7* animals with no mitochondria in the process, we observe the presence of actin dynamics in the neuronal process up to a average distance of ∼78μm from the cell body of the PLM neuron (Fig 3B). The density of actin dynamics near the cell body that includes trails (Fig 3E, F, Table S3) and short-lived actin rich regions (S3A, C, Table S3) are fewer than in wildtype trails (Fig 3E, F, Table S3) and short-lived actin rich regions (S3A, C, Table S3).

Since in *ric-7* animals, there is large number of mitochondria present in the cell body, we addressed the extent of actin dynamics present in *ric-7(lf)* expressing a mTruck transgene that does not restore mitochondria throughout the PLM neuronal process but are present at most 40μm from the cell body along the neuronal process. In these animals we again observe that both actin trails and short-lived actin rich regions are present at a average distance of ∼79 μm from the last mitochondrion along the neuronal process (Fig 3C, D). Beyond these 80 μm after the last mitochondria along the neuronal process, no dynamic actin is observed (Fig 3C).

These data suggest that axonal mitochondria are promote dynamic actin locally. This regulation can extend around 70-80μm from the position of the last mitochondrion.

### Mitochondrial superoxide dismutase, not CED-9, affects actin dynamics

Mitochondria-driven modulation of actomyosin structure in epithelial cells depends on mitochondrial ROS during tissue repair (13, 14). Actin severing in synapses of developing motor neurons *in vivo,* depends on CED-9 (human ortholog of BCL2 like 2 protein)-CED-3 [human ortholog of CASP3 (caspase 3); CASP6 (caspase 6); and CASP7 (caspase 7)] pathway in the context of tissue remodeling (15). To investigate whether these pathways also play a role in the mitochondria-dependent actin dynamics in adult healthy axons *in vivo*, we imaged actin dynamics in loss of function mutants of mitochondrial SOD or CED-9.

We performed time-lapse imaging of GFP::UtCH in loss-of-function mutants of the caspase CED-3 and mitochondrially localized antiapoptotic protein CED-9. Dynamic actin remains unchanged in the single mutants of *ced-9* (Fig. 4 D and S4I, J, Movie 6, Table S4) and *ced-3* (S4D, I, J, Movie 7, Table S4). To assess if the presence of CED-3/CED-9 influences axonal actin dynamics in the absence of axonal mitochondria, we examined the actin rich regions in the double mutants of *ced-9(lf); ric-7(lf)* and *ced-3(lf); ric-7(lf)*. Actin dynamics in these double mutants are similar to *ric-7(lf)* (Fig. 4B, E, S4E, I, J Movie 8, Movie 9, Table S4). Restoring mitochondria only in the TRNs using mTruck in these double mutants restored dynamic actin (Fig. 4C, F, S4F, I, J, Movie 10, Movie 11, Table S4). This data suggests that the CED-3-CED-9 pathway may not play a major role in mitochondria-mediated actin regulation in axons.

**Figure 4:**
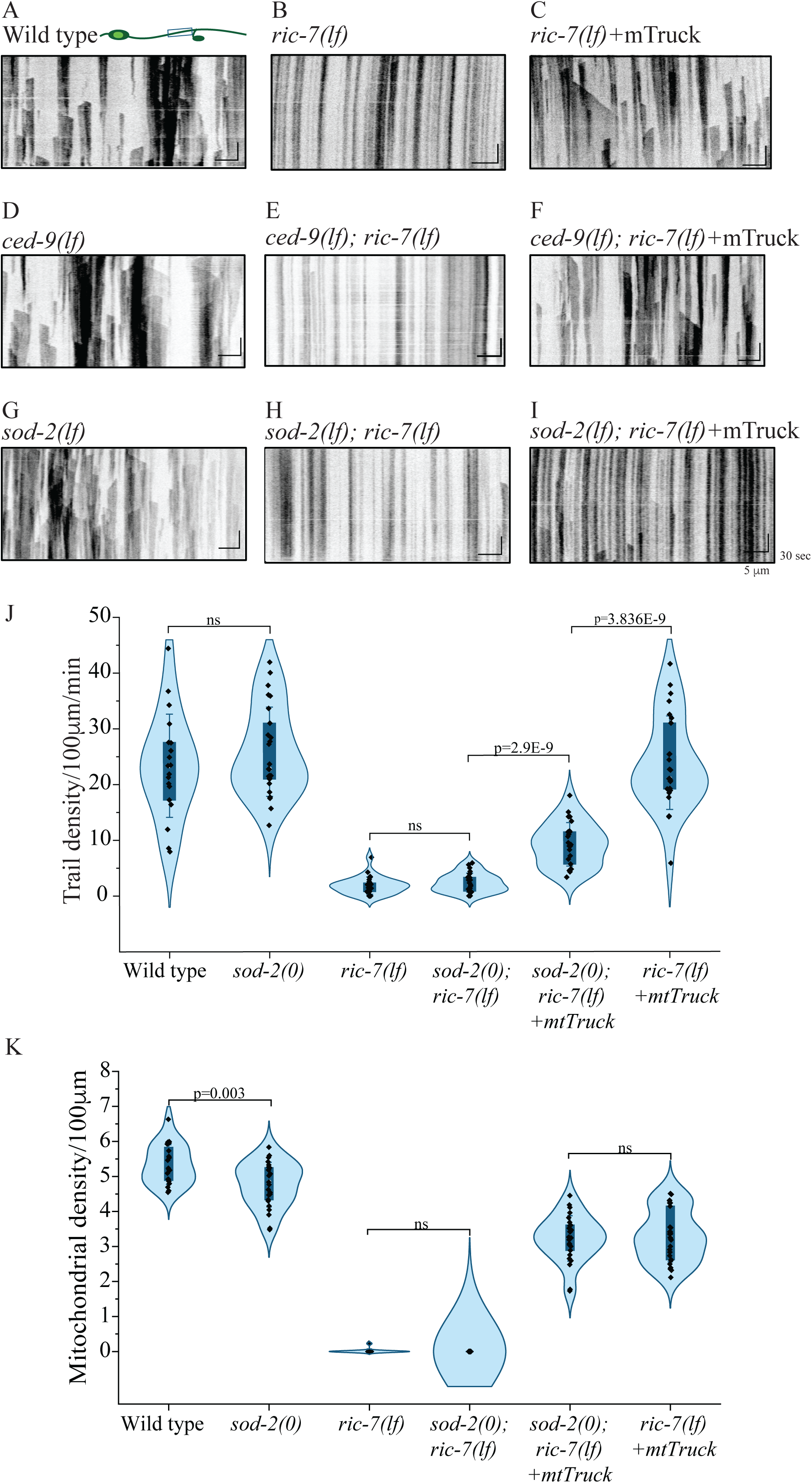
Mitochondria driven actin dynamics are dependent on mitochondrial superoxide dismutase and independent of CED-9. A) Representative kymographs for GFP::UtCH obtained from the PLM major process of wild type. Scale bar: x axis-5 μm, y axis-30 secs. B) Representative kymographs for GFP::UtCH obtained from the PLM major process of *ric-7(nu447).* Scale bar: x axis-5 μm, y axis-30 secs. C) Representative kymographs for GFP::UtCH obtained from the PLM major process of *ric-7(nu447) +tbEx307*(mTruck). Scale bar: x axis-5 μm, y axis-30 secs. D) Representative kymographs for GFP::UtCH obtained from the PLM major process of *ced-9(n2812).* Scale bar: x axis-5 μm, y axis-30 secs. E) Representative kymographs for GFP::UtCH obtained from the PLM major process of *ced-9(n2812); ric-7(nu447).* Scale bar: x axis-5 μm, y axis-30 secs. F) Representative kymographs for GFP::UtCH obtained from the PLM major process of *ced-9(n2812); ric-7(nu447) +tbEx306*(mTruck). Scale bar: x axis-5 μm, y axis-30 secs. G) Representative kymographs for GFP::UtCH obtained from the PLM major process of *sod-2(ok1030)*. Scale bar: x axis-5 μm, y axis-30 secs. H) Representative kymographs for GFP::UtCH obtained from the PLM major process of *sod-2(ok1030); ric-7(nu447).* Scale bar: x axis-5 μm, y axis-30 secs. I) Representative kymographs for GFP::UtCH obtained from the PLM major process of *sod-2(ok1030); ric-7(nu447)* +*ttbEx307*(mTruck). Scale bar: x axis-5 μm, y axis-30 secs. J) Quantitation of trails/100μm/min in *sod-2(ok1030)*, *sod-2(ok1030); ric-7(nu447), sod-2(ok1030); ric-7(nu447)* +*tbEx307*(mTruck). n=25 animals. Two-sample t-test. I) Quantitation of mitochondrial density/100μm in major process of PLM in wild type *sod-2(ok1030)*, *sod-2(ok1030); ric-7(nu447), sod-2(ok1030); ric-7(nu447)* +*tbEx307*(mTruck). n=25 animals. Two-sample t-test.

We examined whether mtROS generated in steady-state mature neurons regulates dynamic actin using the null mutants of *sod-2* (mitochondrial superoxide dismutase). We observe no difference in the actin dynamics between the *sod-2(0)* and wild type (Fig. 4A, G, J, S4G, Movie 12, Table S4). However, restoring mitochondria only in the neuronal process using mTruck in the double mutants of *sod-2(0); ric-7(lf)* failed to fully restore dynamic actin rich regions as compared to *ric-7(lf)*+ mTruck (Fig. 4H, I, J, S4G, Movie 13, Movie 14, Table S4). Although long lived stationary actin rich regions remain unchanged (fig S4H). We examined if the density of mitochondria was altered in genetic backgrounds that contain *sod-2(0)* (Fig. 4K). *sod-2(0); ric-7(lf)* animals continue to lack mitochondria in the axons similar to the single mutant of *ric-7* (Fig. 4K). Likewise, *sod-2(0); ric-7* mutants with TRN-mTruck have mitochondrial densities similar to *ric-7(lf)+* mTruck in the PLM neuronal process (Fig. 4K).

Mitochondria and mtSOD-2 are known to scavenge cytosolic and mitochondrial ROS respectively. Since *sod-2* nulls by themselves do not show changes in actin dynamics but the role for *sod-2* is revealed only in the *ric-7; mTruck* background, we suggest that the reduced mitochondrial density in *ric-7; mTruck* provides a sensitized background where the rolw of ROS in actin dynamics is revealed.

### Release of mitochondrial ROS in the cytosol may regulate dynamic actin in axons *in vivo*

Superoxide dismutase (SOD) quench oxygen anions which are the primary precursors of ROS. *In vitro* studies show that mtSOD-deficient mitochondria release ∼4-fold more superoxides than wild type mitochondria isolated from mouse skeletal muscles (27, 28). Hence, an absence of mtSOD in the axonal mitochondria would likely elevate cytosolic ROS. Likewise, the decreased numbers of mitochondria *in ric-*7 mutants could lead to elevated cytoplamic ROS in neuronal processes. To test this, we used roGFP-tsa2 (29, 30) targeted to the cytosolic side of the plasma membrane using the pleckstrin homology domain PH. roGFP-tsa2 is a ratiometric ROS sensor that allows for normalization of neuronal expression. Since lack of mitochondria along neuronal process abrogates actin dynamics in all sensory and motor neurons, we used an existing reagent expressed in the AVA command interneurons which also have long neuronal processes, analyzing roGFP_tsa2 signaling at the soma and proximal region of AVA∼ 100-150µm from the soma. We investigated cytoplasmic ROS levels in *ric-7* mutants which do not have dynamic actin. We observe that *ric-7(lf)* have higher levels of cytoplasmic ROS as compared to wild type animals in the proximal region of AVA ∼100µm from the soma but not at the somas of AVAs (Fig. S4L, M, N). Thus, mitochondrial ROS released in the cytosol may negatively regulate dynamic actin in the axons under basal conditions.

### Actin dynamics is necessary and sufficient for UNC-9 and UNC-7 gap junction localization in axons

Actin contributes to anchoring, aggregating/clustering and endocytosis of plasma membrane and gap junction proteins in neuronal and non neuronal cells (31-36). We examined the localization of the gap junction innexin proteins UNC-9 and UNC-7 and the mechanosensitive DEG/ENaC channel MEC-4 in the presence and absence of mitochondria along the neuronal process.

We assessed the localization of the MEC-4 subunit of the amiloride of DEG/ENaC channels and observe a decrease in the density of clusters in *ric-7(lf)* animals that lack mitochondria (mean cluster density=6.6/50μm ± 4.1) compared to wild type (mean cluster density =11.8 /50μm ±4.1 Fig 5A, B). As reported earlier, we observe 2-3 UNC-9 clusters along the PLM process near the cell body (the mean size of each cluster ∼1μm in length) and a single punctum at the distal zone (37) Fig. 5C, G, S5B). Mutants in *ric-7(lf)* have a variable number of clusters ranging from 1-5 clusters/PLM neuron near the cell body (Fig. 5D, G) as well as the length covered by the proximal UNC-9 cluster (called proximal zone length) (Fig. S5A), size of each cluster is ∼1μm in length similar to wild type (mean size wild type= 1.03 μm ±0.17; median size for *ric-7(lf)*=1.0 μm) (Fig. S5B). Similar to UNC-9, we observe a change in number of UNC-7 cluster in the absence of mitochondria [*ric-7(lf)*=1-4 cluster/neuron, wild type mean=2 cluster/neuron, Fig 5H, I]. We also assessed the distribution of synaptic vesicle protein, Synaptogyrin::GFP in *ric-7(lf)* mutants and do not observe any gross differences in distribution (Fig. S5C, D).

**Figure 5:**
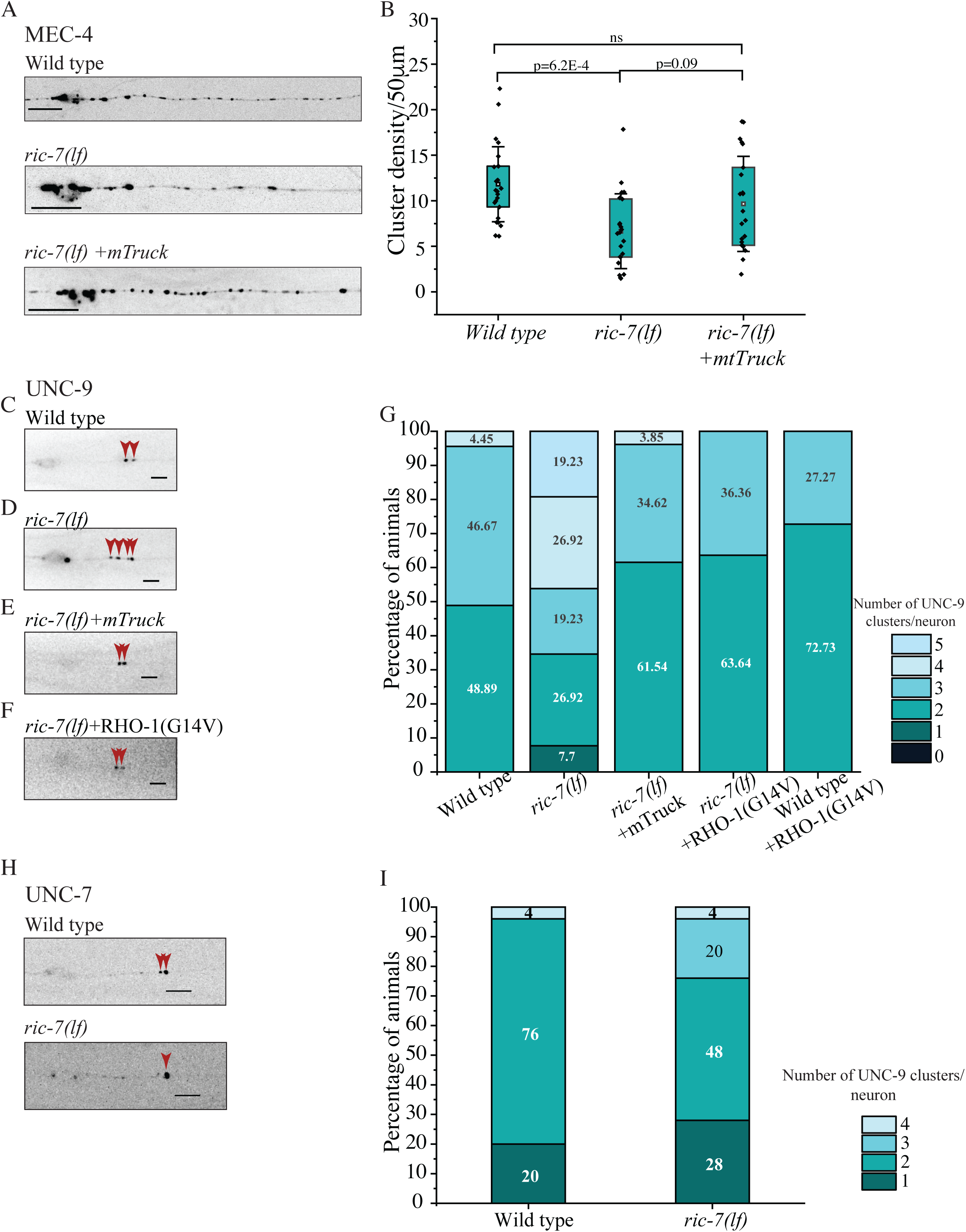
Dynamic actin is important for plasma membrane protein localization in axons. A) Representative fluorescence images of *ljEX437*(MEC-4::mCherry) near the cell body of PLM neuron in wild type, *ric-7(lf)* and *ric-7(lf)*+ mTruck. Scale bar= 10μm B) Quantitation of density of clusters/50μm in wild type, *ric-7(nu447)* and *ric-7(nu447)*+ *tbEx307*(mTruck). n> 20 animals. Kruskal Wallis ANOVA Dunn’s test. C) Representative epifluorescence image of UNC-9::GFP near the cell body of PLM neuron in *tbEx448* (wild type). Scale bar= 5μm red arrohead- UNC-9::GFP cluster. D) Representative epifluorescence image of *tbEx448*(UNC-9::GFP) near the cell body of PLM neuron in *ric-7(nu447).* Scale bar= 5μm. arrohead- UNC-9::GFP cluster. E) Representative epifluorescence image of UNC-9::GFP near the cell body of PLM neuron in *ric-7(nu447)* +*tbEx307*(mTruck). Scale bar= 5μm. arrohead- UNC-9::GFP cluster. F) Representative epifluorescence image of UNC-9::GFP near the cell body of PLM neuron in *ric-7(nu447)* +*twnEx337*[*mec-4*p::RHO-1(G14V)]. scale bar= 5μm. arrohead- UNC-9::GFP cluster. G) Percentage animals with showing distribution of number of clusters of UNC-9::GFP present in wild type, *ric-7(nu447)*, *ric-7(nu447)* +*tbEx307*(mTruck), *ric-7(nu447)* + *twnEx337*[*mec-4*P::RHO-1(G14V)]. H) Representative fluorescence images of *ljEx868*(UNC-7::GFP) near the cell body of PLM neuron in wild type and *ric-7(lf)*. Scale bar= 10μm. arrohead- UNC-7::GFP cluster. I) Distribution of number of clusters of UNC-7::GFP present in wild type and *ric-7(nu447).* n>20 animals

Restoring mitochondria only in TRNs using mTruck in *ric-7* mutants restored the cluster density of MEC-4 (mean cluster density =9.7 /50μm ±5.2) (Fig 5A, B) Similarly, restoring of mitochondria only in TRNs restored these defects in UNC-9 cluster number and proximal zone length (mean cluster size size *ric-7(lf)* +mTruck= 1.08 μm ±0.2) (Fig. 5E, G, S5A). These data suggested that mitochondria and actin dyanamics are necessary for the distribution of UNC-9 and MEC-4.

To assess the role of F-actin in the distribution of these proteins, we rescued the dynamic actin in TRNs alone by expressing constitutively active RHO-1(G14V) in TRNs (Fig S5E). RHO-1(G14V), a mutation that locks RHO-1 in a GTP-bound state (38-40). Prior studies report that RHO-1(G14V) produces a dominant negative effect over endogenous RHO-1(14, 41). We expressed this transgene in TRNs alone and observe that mitochondria continue to be absent in the axons of *ric-7(lf)*+ RHO-1(G14V) however actin dynamics are restored (Fig. S5F). We assessed the localization of UNC-9 in *ric-7(lf)*+ RHO-1(G14V). We observe a rescue in the UNC-9 cluster number and length of proximal region on rescue of dynamic actin (Fig 5F, G, S5A). These data suggests that dynamic actin independent of mitochondria is sufficient for UNC-9 distribution along the neurons.

Thus, mitochondria/mitochondrially driven dynamic actin are necessary and likely sufficient for the distribution of several plasma membrane proteins in TRNs.

### Spectrin organization can occur independent of mitochondria

Periodic spectrin scaffold alternating with the actin rings has been shown in neurons in culture and *in vivo* (3, 5, 42, 43). The MPS has been shown to influence the distribution of plasma membrane proteins (3, 44, 45). We investigated whether dynamic actin that we see alters the distribution of plasma membrane proteins may arise from changes in the MPS. We examined whether absence of mitochondria also affect the lattice periodicity of Spectrin in PLM neurons. We expressed split-GFP reporter system consisting of 7xspGFP11 inserted in the endogenous α-spectrin locus and a spGFP1-10 expressed from cell-specific promoters (43, 46). This system allows the labeling of endogenous spectrin to a single axon. We acquired airyscan images of Spectrin in wildtype and *ric-7* TRNs. Wild type showed showed the presence of periodic Spectrin rings and in *ric-7* animals show a spectrin MPS pattern closer to wildtype than to solulable GFP (Fig S5G, H). These data suggests that Spectrin rings can form independent of dynamic actin in the axons.

### Mitochondria mediated actin dynamics is necessary and sufficient for avoidance behavior in response to gentle touch

We investigated the importance of mitochondrially regulated dynamic actin on the activity of touch receptor neurons *in vivo* by evaluating the escape response after a gentle touch stimulation of the animals. As observed previously, *ric-7(lf)* were defective to gentle touch response (median touch response=60%) [(47), Fig 6A]. This defect was rescued on restoring mitochondria in the neuronal processes of TRNs alone (median touch response=100%) (Fig 6A). These data suggest that mitochondria and mitochondrially regulated actin dynamics are both necessary for gentle touch responsiveness.

**Figure 6:**
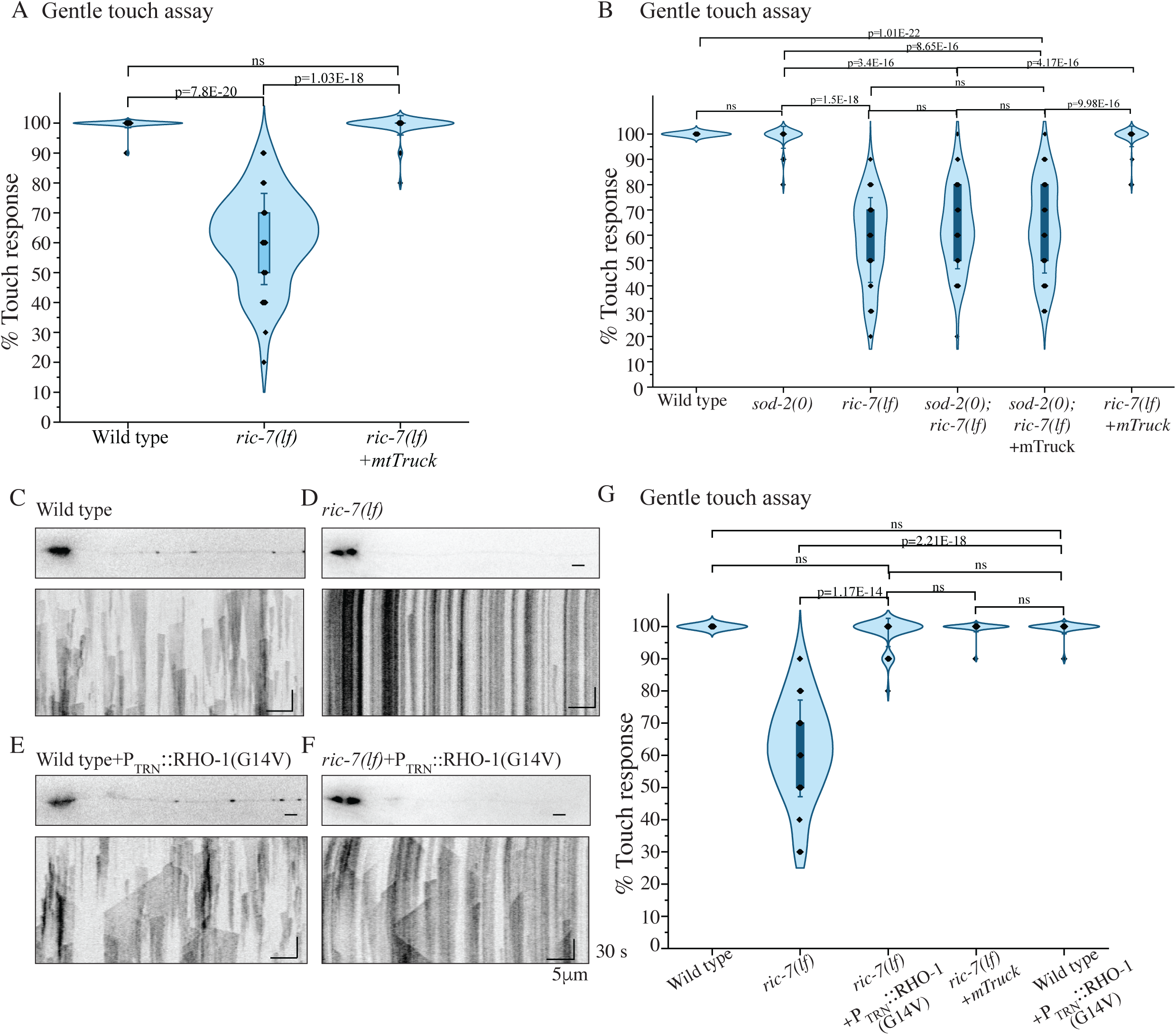
Mitochondria mediated actin dynamics is necessary and sufficient for avoidance behavior in response to gentle touch. A) Percentage touch response in *ric-7(nu447)* and *ric-7(nu447)+ tbEx307*(mTruck). n>50 animals. Kruskal Wallis ANOVA Dunn’s test. B) Percentage touch response in *sod-2(ok1030), sod-2(ok1030); ric-7(nu447)* and *sod-2(ok1030); ric-7(nu447)+ tbEx307*(mTruck). n>50 animals. Kruskal Wallis ANOVA Dunn’s test. C) Top panel: representative mitochondrial images, bottom panel: representative kymographs of GFP::UtCH for wild type. Scale bar: x axis=5μm, y axis=30 sec. D) Top panel: representative mitochondrial images, bottom panel: representative kymographs of GFP::UtCH for *ric-7(nu447).* Scale bar: x axis=5μm, y axis=30 sec. E) Top panel: representative mitochondrial images, bottom panel: representative kymographs of GFP::UtCH for wild type+ *tbIs574*[P_TRN_::RHO-1(G14V)] [P_TRN_: *mec-4p*;TRN specific expression]. Scale bar: x axis=5μm, y axis=30 sec. F) Top panel: representative mitochondrial images, bottom panel: representative kymographs of GFP::UtCH for *ric-7(nu447)*+ *tbIs574*[P_TRN_::RHO-1(G14V)] [P_TRN_: *mec-4*p;TRN specific expression]. Scale bar: x axis=5μm, y axis=30 sec. G) Percentage touch response in wild type, *ric-7(nu447), ric-7(nu447)*+ *tbIs574*[P_TRN_::RHO-1(G14V)] [P_TRN_: *mec-4*p;TRN specific expression], wild type+ *tbIs574*[P_TRN_::RHO-1(G14V)] [P_TRN_: *mec-4*p;TRN specific expression]. n>50 animals. Kruskal Wallis ANOVA with Dunn’s test.

To investigate the contribution of mitochondrially driven dynamic actin, we assessed the gentle touch responsiveness in animals that contain axonal mitochondria but lack dynamic actin due to the absence of *sod-2*. We observe that genotypes such as *sod-2(lf)* which contain both axonal mitochondria and actin dynamics show a touch response similar to wild type animals. *sod-2(lf); ric-7(lf)* that lack axonal mitochondria as well as dynamic actin have a lower touch response (mean touch response=60% ±16.2 S.D) as compared to wild type and are similar to *ric-7(lf)* (median touch response=60%) (Fig 6B), *sod-2(lf); ric-7(lf)+* mTruck which have axonal mitochondria but decreased dynamic actin show a decreased response (median touch response=60%) similar to *ric-7(lf)* (Fig 6B). These data suggested that the presence of actin dynamics but not mitochondria correlates with gentle touch responsiveness in *ric-7* mutants.

We asked if dyanmic actin alone in the absence of mitochondria that restores gap junction localization is sufficient to restore gentle touch responsiveness to *ric-7(0)* animals. We carried out the touch response assay constitutively active RHO-1 and show that these animals had improved touch responsiveness compared to *ric-7*(Fig. 6E, G). Expressing RHO-1 alone in a wild type background showed a touch responsiveness similar to wild type. These data suggest that restoring actin dynamics in the absence of mitochondria is sufficient for touch response in a *ric-7* mutant.

## Discussion

In this study, we report that axonal mitochondria regulate the axonal dynamic actin *in vivo*, possibly through modulating cytosolic ROS levels. We observe that dynamic actin is necessary and sufficient for the escape response of *C. elegans* by regulating the distribution of plasma membrane proteins influencing mechanosensation but not through organization of the MPS.

Dynamic actin can be modulated by multiple regulators in a cell (8-12). Our study shows that axonal mitochondria regulate dynamic actin in neuronal processes *in vivo* (Fig 2 G). Previous studies demonstrate that the dynamic pool of axonal F-actin polymerize from endosomes in vertebrate cultured neurons (1). In contrast, previous data from our lab shows that only 6 % of dynamic actin in axons are associated with stationary pre-synaptic vesicles *in vivo* (2). This suggests that the origin of dynamic actin may vary between neuron types and model systems. Similar to prior studies during tissue remodelling (13, 14), we observe mtROS to be important in regulating dynamic axonal actin in mature neurons *in vivo*. Consistent with a role for ROS, we also see that in *ric-7* animals cytosolic ROS is elevated (Fig S4 P). Elevated ROS is known to oxidize monomeric actin which decreases the rate of actin polymerization and its interaction with actin binding proteins like Profilin, Filamin, etc (48, 49). Elevated ROS fragments and depolymerizes filamentous actin and decreases their presence in the cell (48, 50-52). ROS also differentially affects the actin regulatory proteins- RHO and RAS GTPase by oxidizing their cysteine residues (53, 54). It inactivates RHO GTPase while activates RAS GTPases, indirectly regulating actin treadmilling in the cell (14, 53, 54). Thus changes in cytosolic ROS due to the lack of mitochondria probably acts through multiple pathways to prevent actin dynamics.

Actin forms a physical barrier for lysosomal pausing in dendrites and cargo trafficking in axons (2, 55). It helps distribute plasma membrane proteins and aids in anchoring and maintaining the turnover of gap junction proteins at the plasma membrane (31, 33, 35, 36). Stable axonal actin rings provide mechanical support to axons and supports the function of ion channels along the axon initial segment (AIS) (3, 4, 56). Dynamic actin is proposed to supply F-actin in the pre-synaptic bouton but its contribution in neuron function is poorly understood (1). Assessment of *ric-7(lf)* animals allowed us to investigate the role of dynamic axonal actin in neurons *in vivo*. We show that dynamic axonal actin is necessary and sufficient for the *C. elegans* escape behavior (Fig 6G). The behaviour is a culmination of a) neuron activation on touch b) elevation of intracellular calcium and c) relay of the signal to the post-synaptic neurons. The touch response depends on the ability to sense touch through the mechanically gated DEG/ENaC channels and the innexins UNC-7 hemichannels present in TRNs present in the plasma membrane (57-59). Additionally, gap junction proteins like UNC-9 and potentially UNC-7 in TRNs form electrical synapses with the post-synaptic inter-neurons (60, 61). Prior studies suggest that Actin associates with the connexin/innexin family proteins and can in the absence of actin can alter the localization of connexin/innexins (35, 62). We show that dynamic axonal actin regulates the distribution of plasma membrane proteins implicated in both mechanosensation and in relay of signal to interneurons in the mechanosensory circuit (Fig 5).

Actin in non-neuronal cells is also known to regulate the distribution of plamsa membrane protein by preventing their lateral motion (63-67). Additionally, it aids in both clathrin mediated endocytosis and restricts the movement of clathrin coated vesicles at the plasma membrane (32, 36, 68, 69). Thus, dynamic actin in axons could regulate plasma membrane protein distribution either by regulating their endocytosis. Specifically UNC-9 localization may be regulated by endocytosing these connexin subunits outside the gap junction ensuring a tightly clustered electrical synapse.

## Acknowledgement

We thank Prof William Schafer for providing *mec-4p::*MEC-4::mCherry and *mec-4p::*UNC-7::GFP strains, Prof. Chun-liang Pan for *zdIs5;twnEx337* and plasmid TTpl772, Prof. Anindya Ghosh Roy for UNC-9::GFP strain. We thank Tanushree Pathank for micro-injecting TTpI772, Ritabhas Das for microinjecting TTpI735 and Anusheela Chatterjee for building the strain *miro-1(tm1966);jsIs609*.Some strains were provided by the CGC, which is funded by the NIH Office of Research Infrastructure Programs (P40 OD010440). Research in the Sandhya Koushika’s lab is supported by grants from DAE (1303/2/2019/R&D-II/DAE/2079), PRISM (12-R&D-IMS-5.02-0202), and Howard Hughes Medical Institute International Early Career Scientist Grant 55007425. SY is funded by the NIH grant R35(GM131744), GM133573 and and NS114400. OG is supported by a Walter-Benjamin Scholarship funded by the Deutsche Forschungsgemeinschaft (DFG, German Research Foundation) Project# 465611822. FH and ED were funded by NINDS NS115947.

## Materials and Methods

### Worm maintenance and strains used

Animals were grown on 60 mm plates containing Nematode Growth Medium (NGM) agar media seeded with *E. coli* strain OP50 maintained under standard laboratory conditions at a temperature of 20°C (70). 60 mm plates were obtained from Praveen Scientific (New Delhi, IN), Bacto Agar and Peptone were obtained from BD bioscience (New Jersey, USA), NaCl was obtained from Hi-Media (Mumbai, IN), Cholesterol, CaCl2, MgSO4 from Sigma-Aldrich (St. Louis, Missouri, United States). Strains and transgenes used in the study are described in Table S5.

### Cloning

MitoTruck(TTpI602) was prepared from PTT58 *[unc129p::unc-116::tagRFP::tom7]* plasmid gifted by from Kaplan lab (Dept. of Mol. Biol., Harvard University). *unc-116::tagRFP::tom7* including *unc-54* 3’-UTR was PCR amplified using Phusion polymerase (NEB, Ipswich, MA, USA) from PTT58 *[unc129p::unc-116::tagRFP::tom7]* using following primers, FP: 5’-AGCAAGGCTAGCCAAGACAAGTTTGTAC-3’ and RP: 5’-ACTCACGGGCCCTAGTGGGCAGATCTT-3’ and cloned between Nhe-1 and Apa1 (NEB, Ipswich, MA, USA) sites in TTpl503 *[mec4p::Lamp-1::GFP]*

### Transgenic lines

Transgenic lines were prepared by injecting a cocktail of plasmids in 1d adult N2 strain using using Eppendorf FemtoJet microinjector (Hamburg, Germany) fitted in Olympus IX53 (Tokyo, Japan). Plasmids were purified using Macherey-Nagel NucleoSpin Plasmid purification kit (Düren, Germany) and subjected to Ethanol precipitation before using.

*tbEx307, tbEx306* (TRN specific Mitotruck) A cocktail of three plasmids, 20ng/μl TTpl602 *[mec4p::unc-116::tagRFP::tom7]*, 50 ng/μl TTpl541 *[ttxp::*RFP], 130 ng/μl TTpl542 [PBluscript SK-] was prepared and microinjected into 1d adult N2 strain. For tbEx448, a cocktail of three plasmids 5 ng/μl, TTpI735 [*mec-4p*::UNC-9::GFP] + 10ng/μl, TTpI580[*myo2p*::mCherry] + 185ng/μl TTpI542 [pBluescript SK-] was prepared and microinjected into 1d adult N2 strain.

### Imaging

For all experiments, animals were transferred to a fresh plate at larval stage 4 from a non-contaminated, non-crowded plate and imaged the Posterior Lateral Microtubule neuron (PLM) at young adult stage unless specified. For static imaging, animals were anaesthetized using 1-10 mM sodium azide (obtained from Sigma-Aldrich, St. Louis, Missouri, United States) prepared in M9 buffer. For time-lapse imaging, 5mM tetramisole hydrochloride (obtained from Sigma-Aldrich, St. Louis, Missouri, United States) prepared in M9 buffer was used unless specified unless specified. Animals were anaesthetized on a glass slide(Bluestar No.1 coverslips, Mumbai, IN) with 5% Agarose (obtained from Sigma-Aldrich, St. Louis, Missouri, United States)mounted in a coverslip (Bluestar No.1 coverslips, Mumbai, IN) unless specified. For all datasets, imaging was performed over 3 days.

### Mitochondrial imaging and image analysis

Strains with MLS::GFP fluorophore were used to acquire static epifluorescent mitochondrial images. Images were acquired at 60X/1.35 NA objective using an inverted Olympus IX73 epifluorescence microscope (Tokyo, Japan) equipped with Photokinetics Evolve EMCCD camera with a pixel size of 0.265μm/pixel. Images were taken using the GFP filer, 100% lamp power at an exposure of 150ms and EM gain of 250. Images ending and beginning with overlapping neuronal regions were acquired covering the length of PLM neuron. Overlapping images were then used to reconstruct the entire neuronal process for further quantification. Each slide was imaged with 10 mins of its preparation.

A cut-off of 2×2 pixel was used to identify smaller mitochondria, any fluorescent particle smaller than that was discarded. The total number of mitochondria and the total neuronal length was calculated. Density of mitochondria was calculated by taking a ratio of mitochondrial number by neuronal length. Final representation was done in the form og density of mitochondria/100μm.

### Actin imaging and image analysis

#### a. Hamamatsu spinning disc with Volocity

Time-lapse imaging of GFP::UtCH was performed using Olympus IX83 microscope with Perkin Elmer Ultraview Spinning Disc confocal Yokogawa CSU-X1 module leading to a Hamamatsu EM-CCD camera. Imaging was carried out at 488 nm solid state LASER at 10% LASER power using a 100×/1.4 N.A. oil objective with an effective pixel size of 0.129 μm/pixel. The exposure time was set to 250ms with sensitivity of 169 and a frame rate of 1 frame per second for a period of 3 mins.

#### b. Prime-EM spinning disc with cellsens

GFP::UtCH was imaged on Olympus IX83 microscope with spinning disc fitted with a Yokogawa CSU-W1module leading to a Prime BSI back illuminated sCMOS camera. Imaging was performed using 473 nm solid-state LASER at 5% LASER power, 100×/1.4 NA oil objective with a Prime BSI sCMOS camera configured with 2×2 binning and the SoRa module selected in the cellsens software giving an effective pixel size of 0.13 μm/pixel. Time-lapse movies were taken for 180 seconds with a frame rate of 1 frame per second and exposure of 300 ms.

To analyze GFP::UtCH events, kymographs were generated using imageJ software. Lines were drawn over the kymograph initiating from the position of appearance of the event and ending where the event disappears. Straight lines constitute of stationary events and slant lines constitute of actin trails. Stationary events lasting for from 2-120 secs were considered to be short lived and the rest long lived. Stationary UtCH::GFP events that decrease in size overtime and disappear are considered as shrinking events. Final representation was done in the form of number of event/100μm/min.

Trails initiating from a pre-existing stationary actin event were considered as trails emerging from stationary actin, trails ending at a pre-existing stationary actin event were considered as trails ending from stationary actin. Trails not associated with any actin event were considered as independent trails.

### Actin Mitochondria overlap analaysis

Static images of MLS::TagRFP (for mitochondria) were acquired on Olympus IX83 microscope with spinning disc fitted with a Yokogawa CSU-W1 module leading to a Prime BSI back illuminated sCMOS camera. Imaging was performed using 561 nm solid-state LASER at 5% LASER power, 100×/1.4 NA oil objective with a Prime BSI sCMOS camera configured with 2×2 binning and the SoRa module selected in the cellsens software giving an effective pixel size of 0.13 μm/pixel.The exposure time was set to 250ms. This was followed by acquiring time-lapse images of UtCH::GFP of the same neuron as before.

Imaging was done at the proximal and distal regions to the cell body in the major process of young adult PLM neurons. The proximal region comprised of the initial 50-100μm of major process from the cell body. The distal region consisted of 50-80μm of major process before the branch point. To observe the extent of presence of dynamic actin in *ric-7(lf)* in the proximal region, the cell body was taken in the field of view along with the major process during both mitochondria and actin imaging. To observe the last dynamic actin in incomplete mitochondrial rescue line, the field of view initiated with the last mitochondrion and the corresponding neuronal process was used of UtCH time-lapse imaging.

### Imaging of cytoplasmic ROS

Worms expressing genetically encoded ROS sensor roGFP-tsa2 (see Table for strains, FJH183 and FJH791 both us csfEx61 which contains *flp-18p::PH::roGFP_tsa2*) targeted to the cytoplasmic side of the plasma membrane using the Pleckstrin homology domain PH, in AVA command interneurons, were imaged on an Olympus IX83 microscope with spinning disc fitted with a Yokogawa CSU-X1 module, and Andor iXon ultra EMCCD camera, 405nm and 488nm solid state excitation laser with a 520nm-20 emission filter, and the Metamorph 7.10.1 imaging platform. The quantification of PH::roGFP fluorescence was done using the maximal projection of 21 imaging planes, a region of interest drawn around the neuronal processes yielded the mean fluorescence intensity at 405nm and 488nm excitation. The same region was displaced next to the process for background fluorescence which was then subtracted from the signal before obtaining 405/488 F_ratio_. The same process was performed for fluorescence in the soma of the AVAs except a larger stack of z-planes was used to encompass the entire somatic volumes.

### Airy scan imaging of Spectrin

Nematodes were mounted on a 2% agarose pad and paralyzed in a 5 ul droplet of 10 mM Levamisole (diluted in M9 medium). Images were acquired on an LSM900 inverted microscope (Zeiss), equipped with an Airyscan detector, a plan-apochromat 63x/1.40 oil objective and a 488nm excitation laser. The microscope was operated with Zen blue v. 3.4.91.00000. Raw images were deconvolved in Zeiss blue using standard settings. Deconvolved images were analyzed in Fiji/Image J v2.3.0/1.53f51. To measure lattice periodicity, a 1 pxl (42.5 nm) thick and 2 um long line were drawn in the center of the axonal region to acquire an intensity profile. 3 lines were drawn within each animal to acquire a data point in the autocorrelation amplitude plot. Each intensity profile was used to calculate an autocorrelation function with acf in R. The amplitude of the autocorrelation function was defined as the difference between the first miminum and the subsequent maximum, with the restriction that the maximum had to occur before 425 nm (10pxl).

### Gentle touch assay

Plate preparation: Plates containing NGM media were prepared 4 days before the assay and stored at 4°C were used. Plates were spotted with 400 μl of *E.coli* OP50 (O.D. ∼0.6) one day before the assay and stored at 4°C.

Gentle touch assay: Worms were transferred to fresh touch assay plates at larval stage 4 from a non-contaminated, non-starved and non-crowded plate and assayed at young adult stage (5-6 hours after transfer). The assay was performed in the temperature range of 20-25°C and humidity varied from 50-70%RH. Gentle touch stimulation was provided with the help an eyelash attached to a stick. Worms were touched alternatively just before the pharynx (anterior touch), and anus (posterior touch). A response was counted if worm moved in the direction opposite to the existing direction, accelerated (in case moving along the same direction) or stopped moving. Worms were not touched near the vulval region to avoid an omega turn behavior. The responses were counted using a cell counter wherein, a positive response was counted as 1 and a negative response was counted as 0. After the assay the worm was removed from the plate to avoid assaying it again. The interstimulus interval (ISI) between an anterior and a posterior touch was maintained at 1 second using a counter which beeped after every second.

### Synaptic Vesicle distribution

Strains with SNG-1::GFP fluorophore were used to acquire static epifluorescent images. Images were acquired at 100X/1.40 NA objective using an inverted Olympus IX73 epifluorescence microscope (Tokyo, Japan) equipped with Photokinetics Evolve EMCCD camera with a pixel size of 0.157μm/pixel. Images were taken using the GFP filer, 100% lamp power at an exposure of 500ms and EM gain of 500. Images ending and beginning with overlapping neuronal regions were acquired covering the length of PLM neuron.

### Distribution of gap junction protein UNC-9

GFP labelled UNC-9 were acquired at 60X/1.35 NA objective using an inverted Olympus IX73 epifluorescence microscope (Tokyo, Japan) equipped with Photokinetics Evolve EMCCD camera with a pixel size of 0.265μm/pixel. Images were taken using the GFP filer, 100% lamp power at an exposure of 300ms and EM gain of 300. Images were acquired of young adult PLM neurons were acquired at the cell body, tip of the PLM neuron, ALM cell body and a z stack was taken to capture the nerve ring near the head of the worm.

### Distribution of gap junction protein UNC-7 and MEC-4

Fluorescently tagged UNC-7 (UNC-7::GFP) and MEC-4::mCherry were imaged on Olympus IX83 microscope with spinning disc fitted with a Yokogawa CSU-W1module leading to a Prime BSI back illuminated sCMOS camera. Imaging was performed using 473 nm solid-state LASER at 1% and 561nm at 10% LASER power for UNC-7 and MEAC-4 respectively, 100×/1.4 NA oil objective with a Prime BSI sCMOS camera configured with 2×2 binning and the SoRa module selected in the cellsens software giving an effective pixel size of 0.13 μm/pixel. Time-lapse movies with exposure was set at 300ms with 3 frames per second. The images spanned the entire length of PLM.

### Microtubule polarity

Time lapse imaging of EBP-2::GFP was performed at 60X/1.35 NA objective using an inverted Olympus IX73 epifluorescence microscope (Tokyo, Japan) equipped with Photokinetics Evolve EMCCD camera. Imaging was carried out using GFP filer, 100% lamp power at an exposure of 400ms and EM gain of 300 with 2 frames per second for 2 mins. Major and minor processes of the young adult PLM neuron were imaged.

### Statistical tests

Shapiro–Wilk test was used on each sample set to test the normality of the distribution. Welch’s t-test was used to compare the means of distributions that passed the test of normality but differ from each other in their variance and sample sizes. Leven’s test to check for equal variance between groups to be compared. Most data is plotted as a violin plot, box representing the 25^th^-75^th^ percentile, mean/median and individual datapoints overlapping over the box plot unless specified. Whiskers represent ± SD. All the data was plotted using OriginPro 2020b (Origin Lab, Northampton, MA, USA). Figures are prepared using Adobe Illustrator (Adobe Corporation, San Jose, CA, USA).

## Reference

1. A. Ganguly et al., A dynamic formin-dependent deep F-actin network in axons. J Cell Biol 210, 401–417 (2015).

2. P. Sood et al., Cargo crowding at actin-rich regions along axons causes local traffic jams. Traffic 19, 166–181 (2018).

3. K. Xu, G. Zhong, X. Zhuang, Actin, spectrin, and associated proteins form a periodic cytoskeletal structure in axons. Science 339, 452–456 (2013).

4. C. Leterrier et al., Nanoscale Architecture of the Axon Initial Segment Reveals an Organized and Robust Scaffold. Cell Rep 13, 2781–2793 (2015).

5. J. Pielage et al., A presynaptic giant ankyrin stabilizes the NMJ through regulation of presynaptic microtubules and transsynaptic cell adhesion. Neuron 58, 195–209 (2008).

6. Y. Qu, I. Hahn, S. E. Webb, S. P. Pearce, A. Prokop, Periodic actin structures in neuronal axons are required to maintain microtubules. Mol Biol Cell 28, 296–308 (2017).

7. S. Dubey et al., The axonal actin-spectrin lattice acts as a tension buffering shock absorber. Elife 9 (2020).

8. B. W. Bernstein, J. R. Bamburg, ADF/cofilin: a functional node in cell biology. Trends Cell Biol 20, 187–195 (2010).

9. P. Lappalainen, T. Kotila, A. Jégou, G. Romet-Lemonne, Biochemical and mechanical regulation of actin dynamics. Nat Rev Mol Cell Biol 23, 836–852 (2022).

10. K. Rottner, T. E. Stradal, Actin dynamics and turnover in cell motility. Curr Opin Cell Biol 23, 569–578 (2011).

11. S. H. Lee, R. Dominguez, Regulation of actin cytoskeleton dynamics in cells. Mol Cells 29, 311–325 (2010).

12. R. Levayer, T. Lecuit, Biomechanical regulation of contractility: spatial control and dynamics. Trends Cell Biol 22, 61–81 (2012).

13. S. Muliyil, M. Narasimha, Mitochondrial ROS regulates cytoskeletal and mitochondrial remodeling to tune cell and tissue dynamics in a model for wound healing. Dev Cell 28, 239–252 (2014).

14. S. Xu, A. D. Chisholm, C. elegans epidermal wounding induces a mitochondrial ROS burst that promotes wound repair. Dev Cell 31, 48–60 (2014).

15. L. Meng et al., The Cell Death Pathway Regulates Synapse Elimination through Cleavage of Gelsolin in Caenorhabditis elegans Neurons. Cell Rep 11, 1737–1748 (2015).

16. A. Ketschek, G. Gallo, Nerve growth factor induces axonal filopodia through localized microdomains of phosphoinositide 3-kinase activity that drive the formation of cytoskeletal precursors to filopodia. J Neurosci 30, 12185–12197 (2010).

17. C. W. Lee, H. B. Peng, The function of mitochondria in presynaptic development at the neuromuscular junction. Mol Biol Cell 19, 150–158 (2008).

18. R. Chakrabarti et al., Mitochondrial dysfunction triggers actin polymerization necessary for rapid glycolytic activation. J Cell Biol 221 (2022).

19. W. K. Ji, A. L. Hatch, R. A. Merrill, S. Strack, H. N. Higgs, Actin filaments target the oligomeric maturation of the dynamin GTPase Drp1 to mitochondrial fission sites. Elife 4, e11553 (2015).

20. A. Gutnick, M. R. Banghart, E. R. West, T. L. Schwarz, The light-sensitive dimerizer zapalog reveals distinct modes of immobilization for axonal mitochondria. Nat Cell Biol 21, 768–777 (2019).

21. K. Tanaka, Y. Sugiura, R. Ichishita, K. Mihara, T. Oka, KLP6: a newly identified kinesin that regulates the morphology and transport of mitochondria in neuronal cells. J Cell Sci 124, 2457–2465 (2011).

22. R. S. Stowers, L. J. Megeath, J. Gorska-Andrzejak, I. A. Meinertzhagen, T. L. Schwarz, Axonal transport of mitochondria to synapses depends on milton, a novel Drosophila protein. Neuron 36, 1063–1077 (2002).

23. X. Guo et al., The GTPase dMiro is required for axonal transport of mitochondria to Drosophila synapses. Neuron 47, 379–393 (2005).

24. Q. Cai, C. Gerwin, Z. H. Sheng, Syntabulin-mediated anterograde transport of mitochondria along neuronal processes. J Cell Biol 170, 959–969 (2005).

25. Y. Wu et al., Polarized localization of kinesin-1 and RIC-7 drives axonal mitochondria anterograde transport. J Cell Biol 223 (2024).

26. R. L. Rawson et al., Axons degenerate in the absence of mitochondria in C. elegans. Curr Biol 24, 760–765 (2014).

27. M. S. Lustgarten et al., Conditional knockout of Mn-SOD targeted to type IIB skeletal muscle fibers increases oxidative stress and is sufficient to alter aerobic exercise capacity. Am J Physiol Cell Physiol 297, C1520–1532 (2009).

28. M. S. Lustgarten et al., Complex I generated, mitochondrial matrix-directed superoxide is released from the mitochondria through voltage dependent anion channels. Biochem Biophys Res Commun 422, 515–521 (2012).

29. R. L. Doser, K. M. Knight, E. W. Deihl, F. J. Hoerndli, Activity-dependent mitochondrial ROS signaling regulates recruitment of glutamate receptors to synapses. Elife 13 (2024).

30. S. De Henau, M. Pagès-Gallego, W. J. Pannekoek, T. B. Dansen, Mitochondria-Derived H(2)O(2) Promotes Symmetry Breaking of the C. elegans Zygote. Dev Cell 53, 263–271.e266 (2020).

31. G. Tadvalkar, P. Pinto da Silva, In vitro, rapid assembly of gap junctions is induced by cytoskeleton disruptors. J Cell Biol 96, 1279–1287 (1983).

32. K. Jordan, R. Chodock, A. R. Hand, D. W. Laird, The origin of annular junctions: a mechanism of gap junction internalization. J Cell Sci 114, 763–773 (2001).

33. G. Gaietta et al., Multicolor and electron microscopic imaging of connexin trafficking. Science 296, 503–507 (2002).

34. R. G. Johnson et al., Gap junctions assemble in the presence of cytoskeletal inhibitors, but enhanced assembly requires microtubules. Exp Cell Res 275, 67–80 (2002).

35. L. Meng, D. Yan, NLR-1/CASPR Anchors F-Actin to Promote Gap Junction Formation. Dev Cell 55, 574–587.e573 (2020).

36. F. Wernert et al., The actin-spectrin submembrane scaffold restricts endocytosis along proximal axons. bioRxiv, 2023.2012.2019.572337 (2023).

37. L. Meng, C. H. Chen, D. Yan, Regulation of Gap Junction Dynamics by UNC-44/ankyrin and UNC-33/CRMP through VAB-8 in C. elegans Neurons. PLoS Genet 12, e1005948 (2016).

38. S. Chen, Z. Zhang, Y. Zhang, T. Choi, Y. Zhao, Activation Mechanism of RhoA Caused by Constitutively Activating Mutations G14V and Q63L. Int J Mol Sci 23 (2022).

39. A. Kumawat, S. Chakrabarty, K. Kulkarni, Nucleotide Dependent Switching in Rho GTPase: Conformational Heterogeneity and Competing Molecular Interactions. Sci Rep 7, 45829 (2017).

40. K. Ihara et al., Crystal structure of human RhoA in a dominantly active form complexed with a GTP analogue. J Biol Chem 273, 9656–9666 (1998).

41. H. A. Benink, W. M. Bement, Concentric zones of active RhoA and Cdc42 around single cell wounds. J Cell Biol 168, 429–439 (2005).

42. G. Zhong et al., Developmental mechanism of the periodic membrane skeleton in axons. Elife 3 (2014).

43. O. Glomb et al., A kinesin-1 adaptor complex controls bimodal slow axonal transport of spectrin in Caenorhabditis elegans. Dev Cell 58, 1847–1863.e1812 (2023).

44. J. Rentsch et al., Sub-membrane actin rings compartmentalize the plasma membrane. J Cell Biol 223 (2024).

45. D. Albrecht et al., Nanoscopic compartmentalization of membrane protein motion at the axon initial segment. J Cell Biol 215, 37–46 (2016).

46. R. Jia et al., Spectrin-based membrane skeleton supports ciliogenesis. PLoS Biol 17, e3000369 (2019).

47. A. Awasthi et al., Regulated distribution of mitochondria in touch receptor neurons of *C. elegans* influences touch response. bioRxiv, 2020.2007.2026.221523 (2020).

48. I. DalleDonne, A. Milzani, R. Colombo, H2O2-treated actin: assembly and polymer interactions with cross-linking proteins. Biophys J 69, 2710–2719 (1995).

49. I. Lassing et al., Molecular and structural basis for redox regulation of beta-actin. J Mol Biol 370, 331–348 (2007).

50. V. Munnamalai, D. M. Suter, Reactive oxygen species regulate F-actin dynamics in neuronal growth cones and neurite outgrowth. J Neurochem 108, 644–661 (2009).

51. J. Sakai et al., Reactive oxygen species-induced actin glutathionylation controls actin dynamics in neutrophils. Immunity 37, 1037–1049 (2012).

52. R. P. Kommaddi et al., Glutaredoxin1 Diminishes Amyloid Beta-Mediated Oxidation of F-Actin and Reverses Cognitive Deficits in an Alzheimer’s Disease Mouse Model. Antioxid Redox Signal 31, 1321–1338 (2019).

53. L. Mitchell, G. A. Hobbs, A. Aghajanian, S. L. Campbell, Redox regulation of Ras and Rho GTPases: mechanism and function. Antioxid Redox Signal 18, 250–258 (2013).

54. J. Heo, K. W. Raines, V. Mocanu, S. L. Campbell, Redox regulation of RhoA. Biochemistry 45, 14481–14489 (2006).

55. B. van Bommel, A. Konietzny, O. Kobler, J. Bär, M. Mikhaylova, F-actin patches associated with glutamatergic synapses control positioning of dendritic lysosomes. Embo j 38, e101183 (2019).

56. S. Vassilopoulos, S. Gibaud, A. Jimenez, G. Caillol, C. Leterrier, Ultrastructure of the axonal periodic scaffold reveals a braid-like organization of actin rings. Nat Commun 10, 5803 (2019).

57. D. S. Walker, W. R. Schafer, Distinct roles for innexin gap junctions and hemichannels in mechanosensation. Elife 9 (2020).

58. N. Tavernarakis, M. Driscoll, Mechanotransduction in Caenorhabditis elegans: the role of DEG/ENaC ion channels. Cell Biochem Biophys 35, 1–18 (2001).

59. S. L. Geffeney et al., DEG/ENaC but not TRP channels are the major mechanoelectrical transduction channels in a C. elegans nociceptor. Neuron 71, 845–857 (2011).

60. M. Chalfie, J. Sulston, Developmental genetics of the mechanosensory neurons of Caenorhabditis elegans. Dev Biol 82, 358–370 (1981).

61. M. Chalfie et al., The neural circuit for touch sensitivity in Caenorhabditis elegans. J Neurosci 5, 956–964 (1985).

62. C. Qu, P. Gardner, I. Schrijver, The role of the cytoskeleton in the formation of gap junctions by Connexin 30. Exp Cell Res 315, 1683–1692 (2009).

63. B. Winckler, P. Forscher, I. Mellman, A diffusion barrier maintains distribution of membrane proteins in polarized neurons. Nature 397, 698–701 (1999).

64. E. S. Wu, D. W. Tank, W. W. Webb, Unconstrained lateral diffusion of concanavalin A receptors on bulbous lymphocytes. Proc Natl Acad Sci U S A 79, 4962–4966 (1982).

65. J. H. Li et al., Directed manipulation of membrane proteins by fluorescent magnetic nanoparticles. Nat Commun 11, 4259 (2020).

66. M. S. Paller, Lateral mobility of Na,K-ATPase and membrane lipids in renal cells. Importance of cytoskeletal integrity. J Membr Biol 142, 127–135 (1994).

67. K. Xu, H. P. Babcock, X. Zhuang, Dual-objective STORM reveals three-dimensional filament organization in the actin cytoskeleton. Nat Methods 9, 185–188 (2012).

68. I. Gaidarov, F. Santini, R. A. Warren, J. H. Keen, Spatial control of coated-pit dynamics in living cells. Nat Cell Biol 1, 1–7 (1999).

69. C. Lamaze, L. M. Fujimoto, H. L. Yin, S. L. Schmid, The actin cytoskeleton is required for receptor-mediated endocytosis in mammalian cells. J Biol Chem 272, 20332–20335 (1997).

70. S. Brenner, The genetics of Caenorhabditis elegans. Genetics 77, 71–94 (1974).

71. P. H. Chia, B. Chen, P. Li, M. K. Rosen, K. Shen, Local F-actin network links synapse formation and axon branching. Cell 156, 208–220 (2014).

72. Y. Hao, Z. Hu, D. Sieburth, J. M. Kaplan, RIC-7 promotes neuropeptide secretion. PLoS Genet 8, e1002464 (2012).

73. Anonymous, large-scale screening for targeted knockouts in the Caenorhabditis elegans genome. G3 (Bethesda) 2, 1415-1425 (2012).

74. G. R. Sure et al., UNC-16/JIP3 and UNC-76/FEZ1 limit the density of mitochondria in C. elegans neurons by maintaining the balance of anterograde and retrograde mitochondrial transport. Sci Rep 8, 8938 (2018).

75. C. Fatouros et al., Inhibition of tau aggregation in a novel Caenorhabditis elegans model of tauopathy mitigates proteotoxicity. Hum Mol Genet 21, 3587–3603 (2012).

76. Q. Zheng et al., The vesicle protein SAM-4 regulates the processivity of synaptic vesicle transport. PLoS Genet 10, e1004644 (2014).

77. C. H. Chen, C. W. He, C. P. Liao, C. L. Pan, A Wnt-planar polarity pathway instructs neurite branching by restricting F-actin assembly through endosomal signaling. PLoS Genet 13, e1006720 (2017).

78. M. Chuang et al., The microtubule minus-end-binding protein patronin/PTRN-1 is required for axon regeneration in C. elegans. Cell Rep 9, 874–883 (2014).

79. S. S. P. Nadiminti, et al., Active zone protein SYD-2/Liprin-α acts downstream of LRK-1/LRRK2 to regulate polarized trafficking of synaptic vesicle precursors through clathrin adaptor protein complexes. bioRxiv (2023).

80. Y. Cho et al., Automated and controlled mechanical stimulation and functional imaging in vivo in C. elegans. Lab Chip 17, 2609–2618 (2017).

81. T. Zhao, Y. Hao, J. M. Kaplan, Axonal Mitochondria Modulate Neuropeptide Secretion Through the Hypoxic Stress Response in Caenorhabditis elegans. Genetics 210, 275–285 (2018).

