## Supplementary material for "Axonal mitochondria regulate gentle touch response through control of axonal actin dynamics"

Supplementary Figure S1

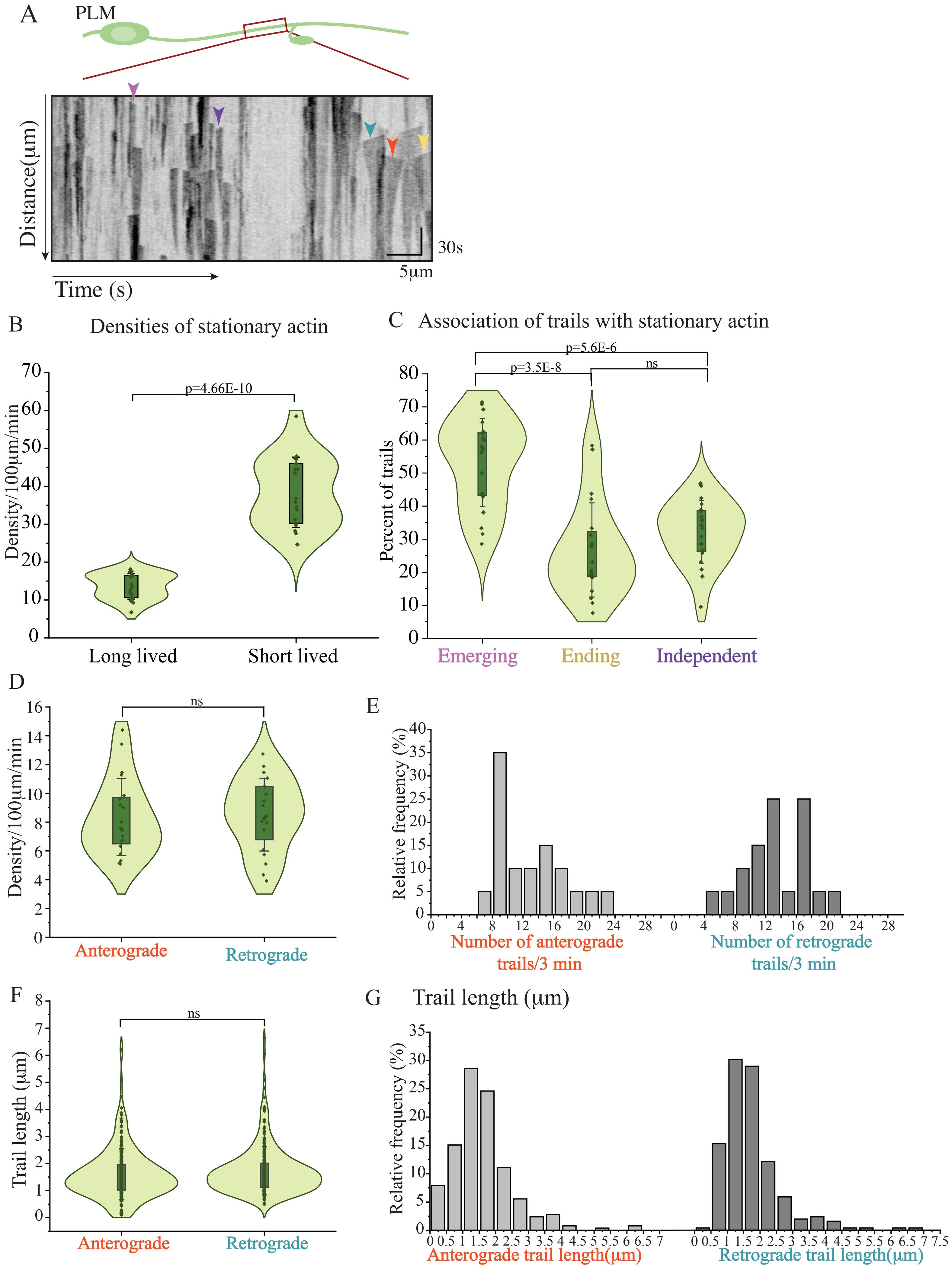

Supplementary Figure S1

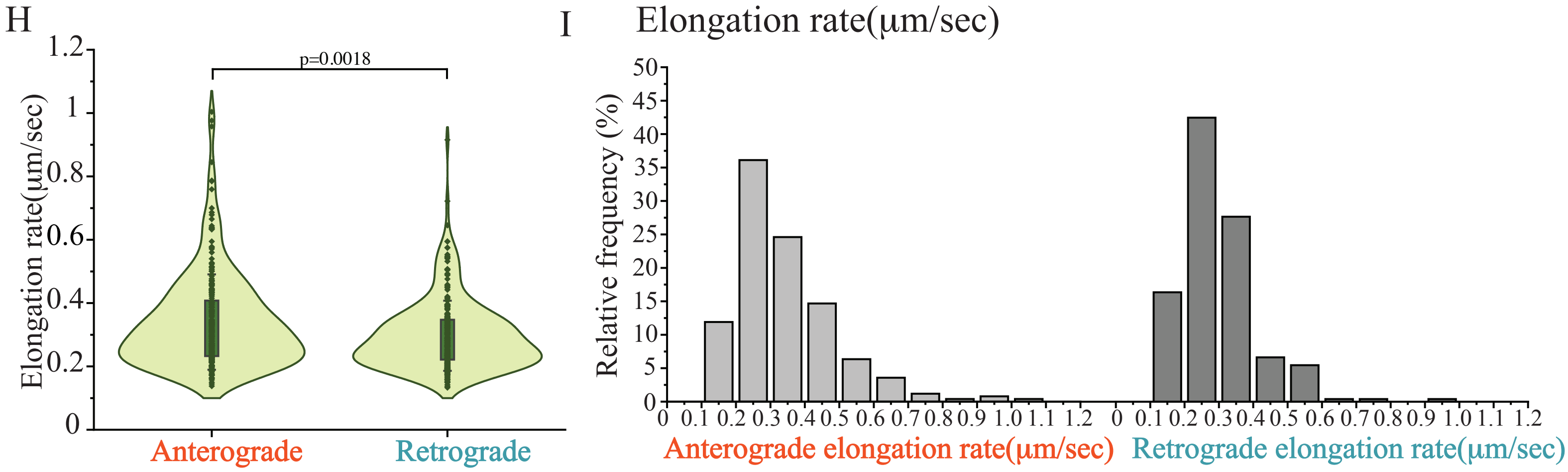

Supplementary Figure S2

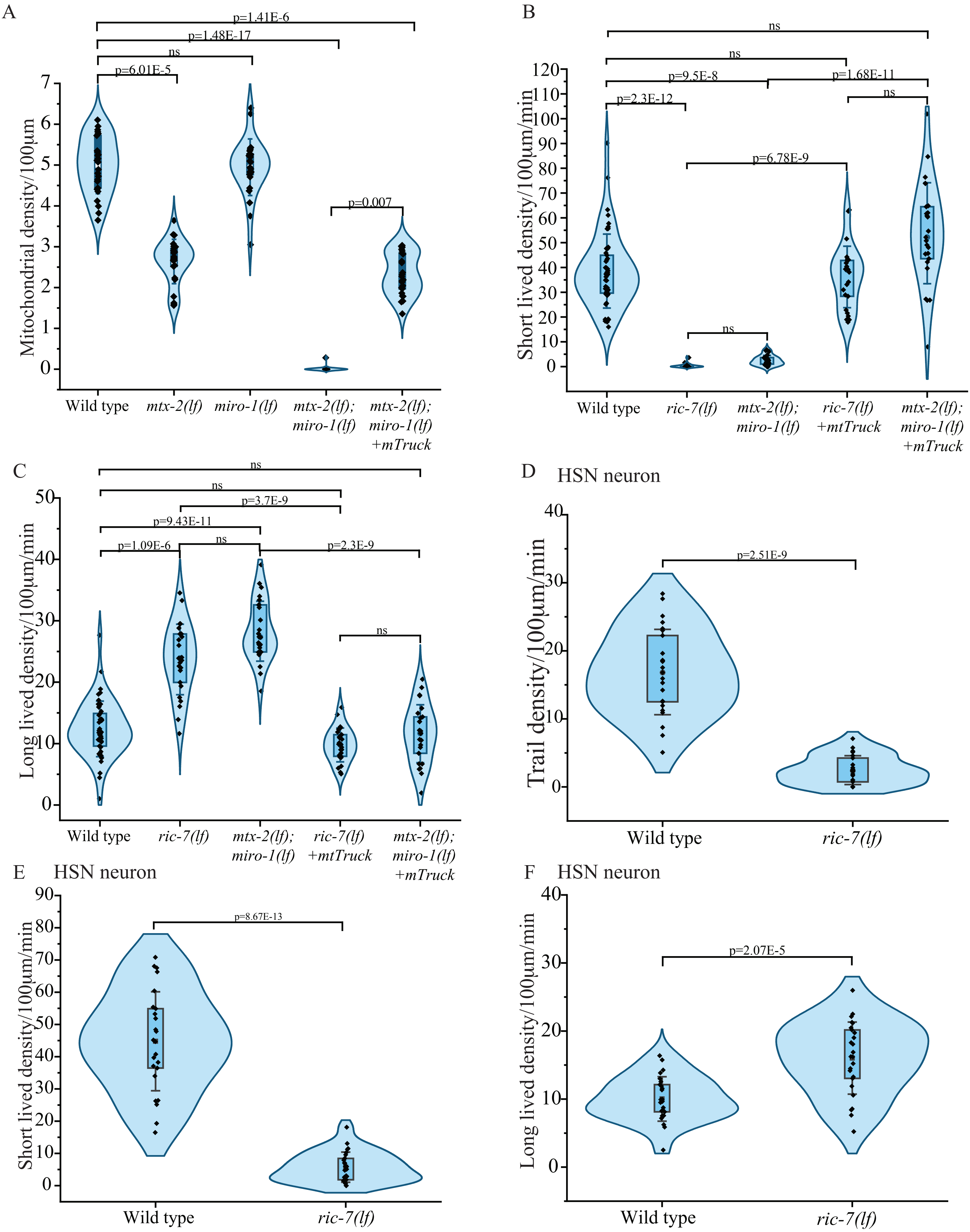

Supplementary Figure S2

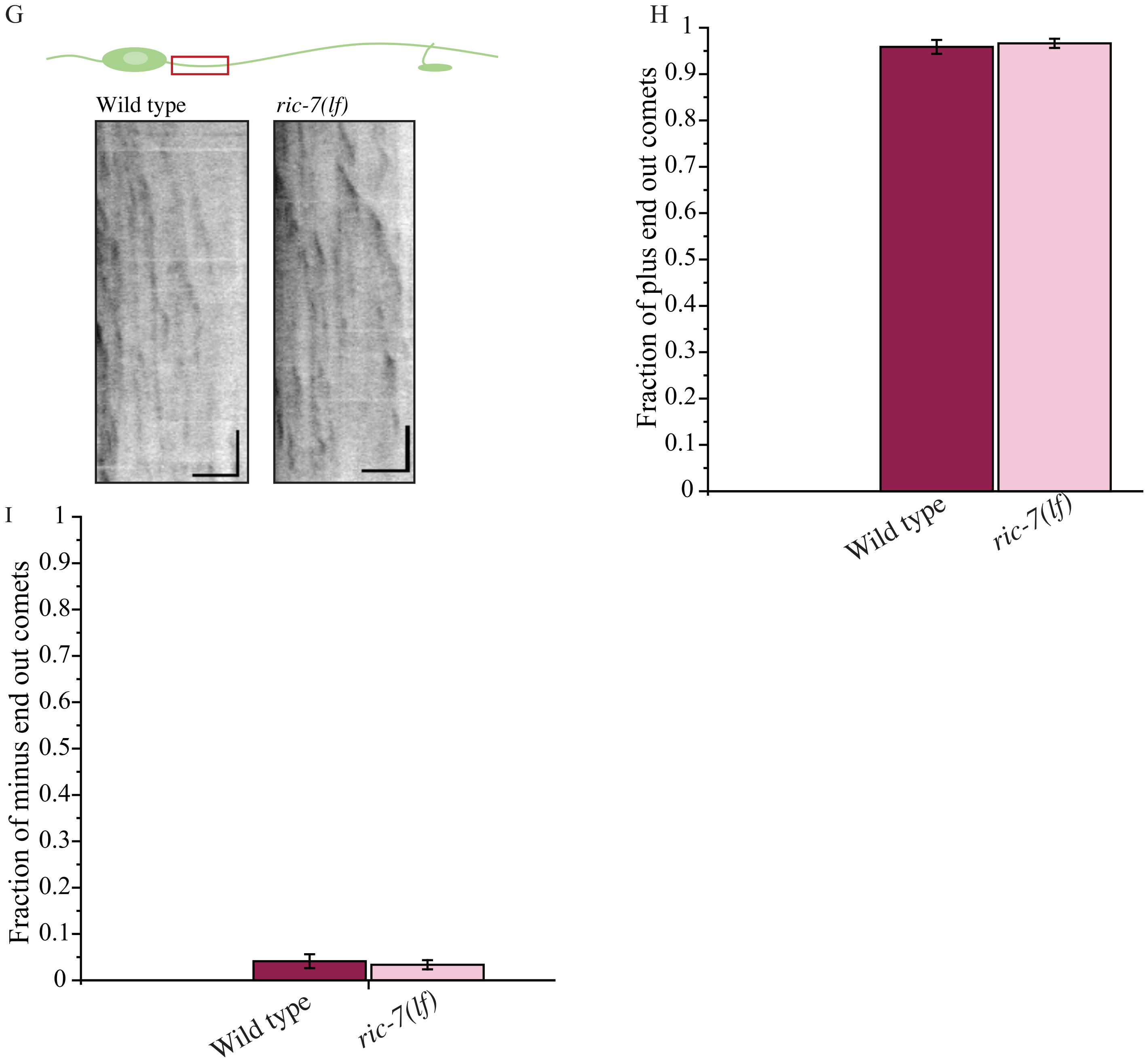

Supplementary Figure S3

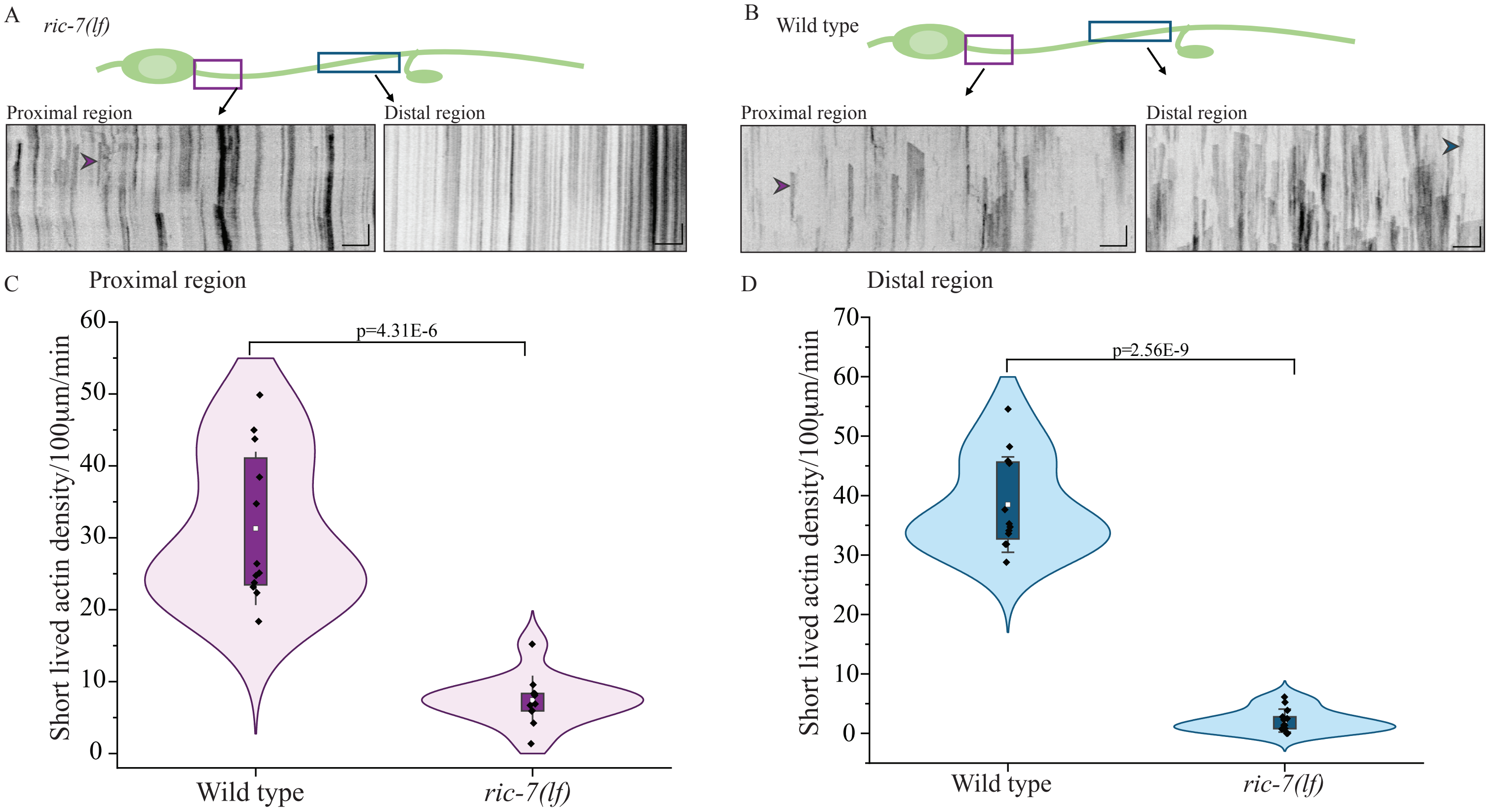

Supplementary Figure S4

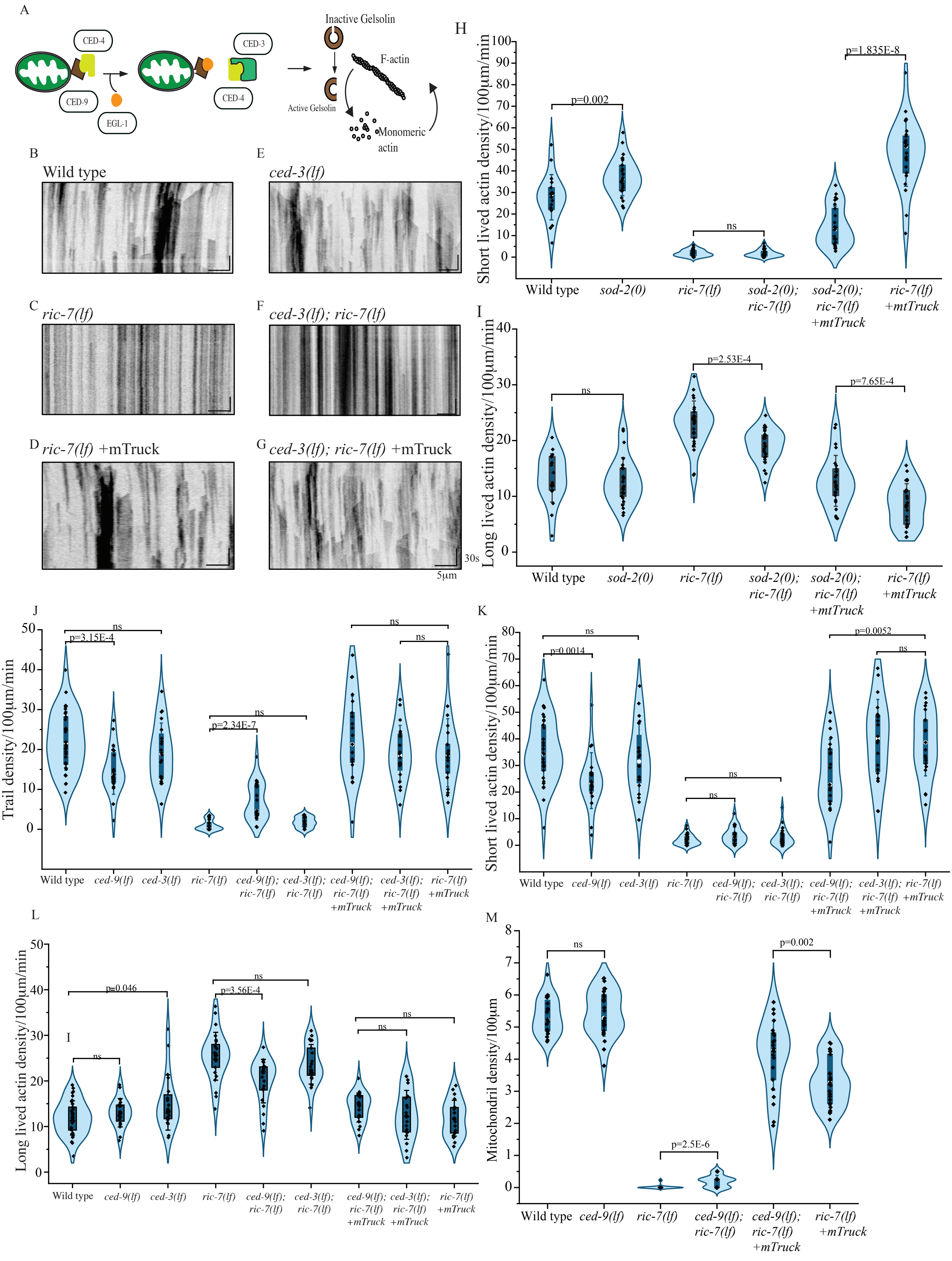

Supplementary Figure S4

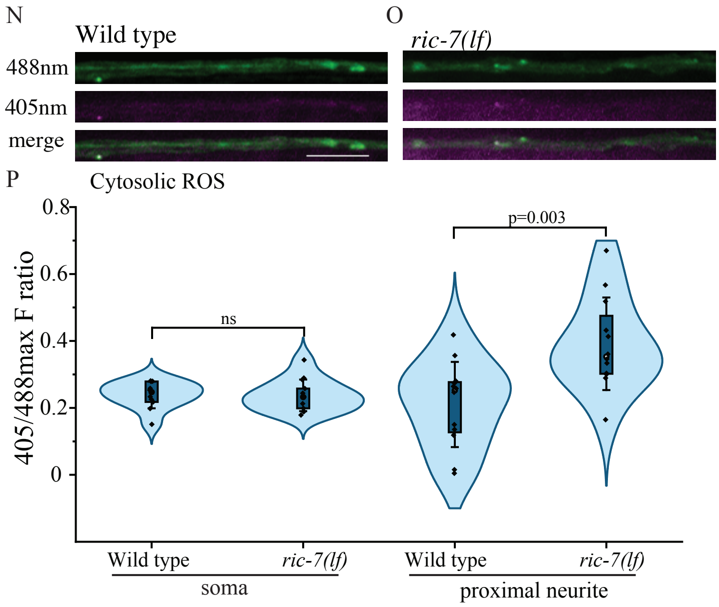

Supplementary Figure S5

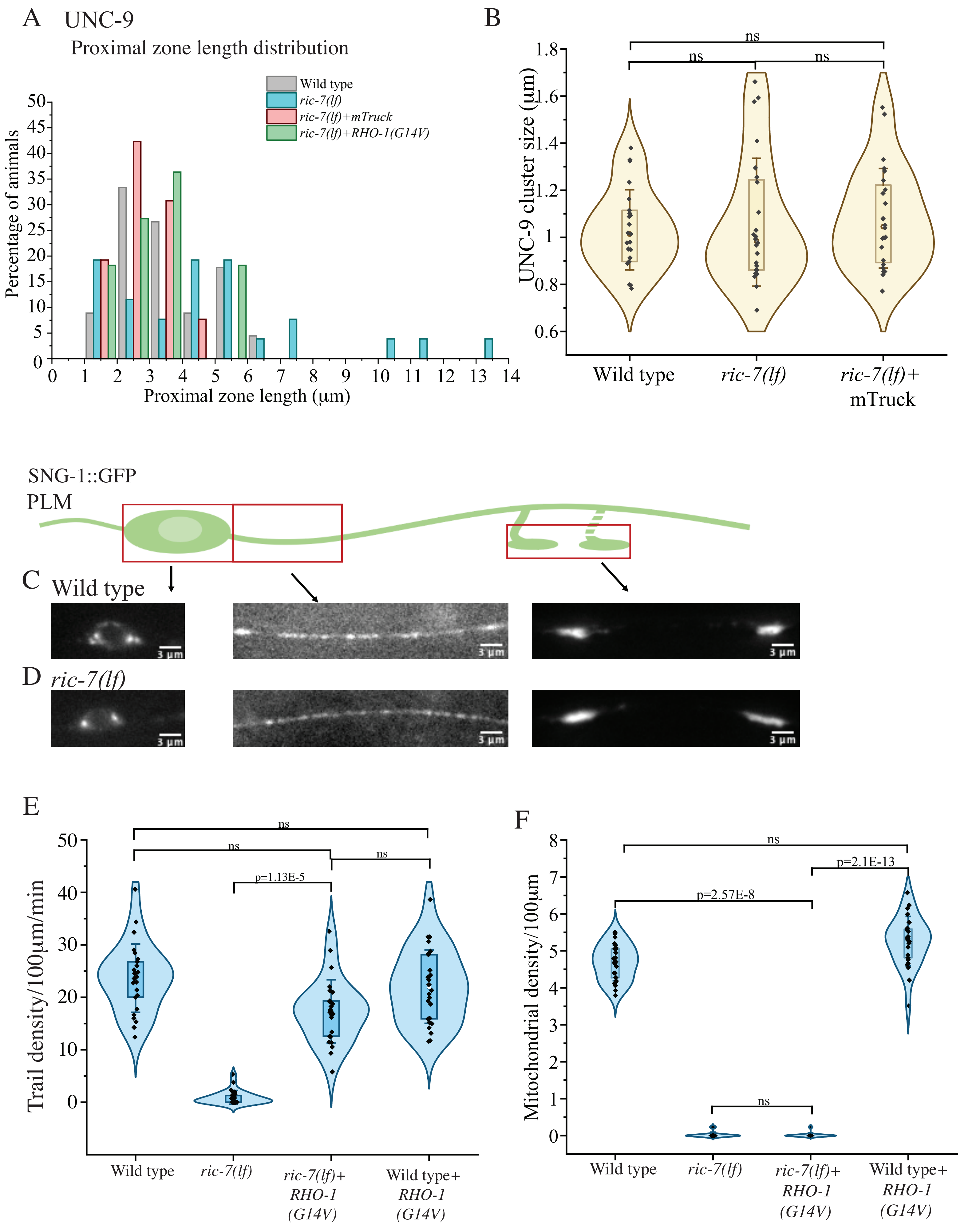

Supplementary Figure S5

G

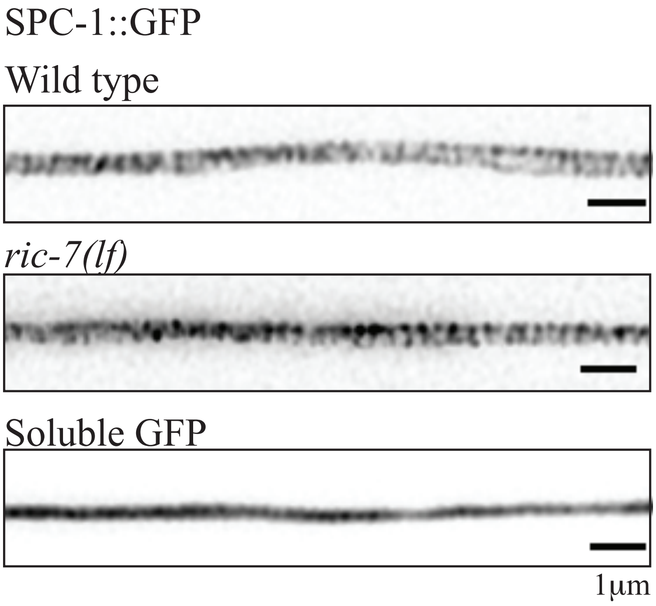


H

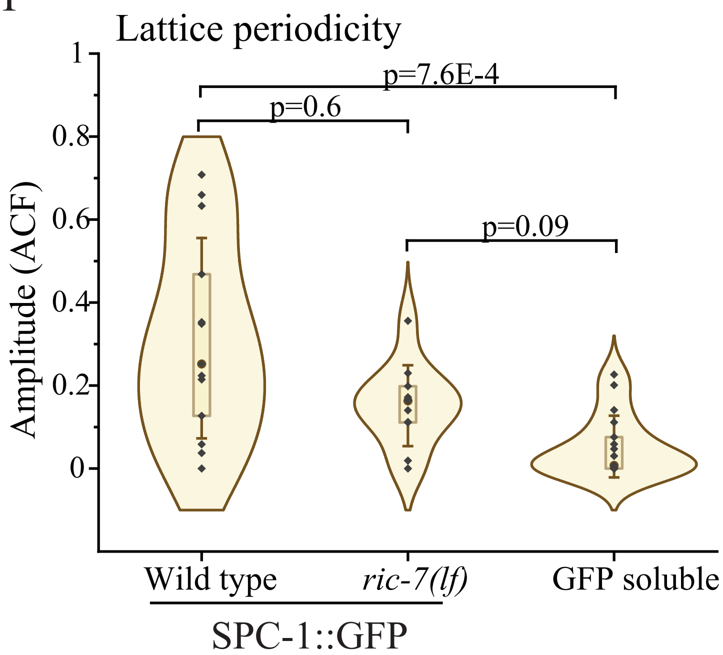

### Supplementary figure legends

#### Supplementary Figure 1:

A) Kymograph obtained by time-lapse imaging of UtCh::GFP in PLM neuron. Arrow heads: Pink- trail emerging from stationary actin, yellow- trail ending in stationary actin, purple- trail independent stationary actin, orange- anterograde trail, cyan- retrograde trail

B) Quantitation of density of short lived and long-lived stationary actin in wild type PLM neuronal process. n=18 animals. Two-Sample t-test with unequal variance, Welch correction.

C) Quantitation of percentage of trails emerging from, ending into and are independent of stationary actin. n=20 animals. One way ANOVA with Bonferroni correction.

D) Quantitation of density of anterograde and retrograde trails. n=20 animals. Two-Sample t-test with equal variance.

E) Relative frequency of anterograde and retrograde trails for 3 mins. n=20 animals.

F) Length (µm) of anterograde and retrograde trails n=252 trails for anterograde trails and 256

trails for retrograde trails. N=20 animals. Mann-Whitney test.

G) Relative frequency (%) of trail lengths in µm n=510 trails. N=20 animals.

H) Elongation rates of anterograde and retrograde trails. n=252 trails for anterograde trails and 256 trails for retrograde trails. N=20 animals. Mann-Whitney test.

I) Relative frequency of elongation rates of anterograde and retrograde trails in 3 mins n=252 trails for anterograde trails and 256 trails for retrograde trails. N=20 animals. Mann-Whitney test.

#### Supplementary Figure 2:

A) Quantitation of mitochondrial density/100µm in major process of PLM in wild type, *mtx-2(gk444), miro-1(tm1966), mtx-2(gk444);miro-1(tm1966), mtx-2(gk444); miro-1(tm1966)+ tbEx307*(mTruck). n≥ 25 animals, Kruskal Wallis ANOVA Dunn's test.

B) Quantitation of short-lived actin density/100µm/min in major process of PLM in wild type, *ric-7(nu447)*, *mtx-2(gk444); miro-1(tm1966)*, *ric-7(nu447) +tbEx307*(mTruck), *mtx-2(gk444); miro-1(tm1966)+ tbEx307*(mTruck). n ≥25 animals for all genotypes, ≥ 80 short-lived actin for *ric-7(nu447)* and *mtx-2(gk444);miro-1(tm1966)*; ≥ 1600 short-lived actin for other genotypes. Kruskal Wallis ANOVA Dunn's test.

C) Quantitation of long-lived actin density/100µm/min in major process of PLM in wild type, *ric-7(nu447)*, *mtx-2(gk444); miro-1(tm1966)*, *ric-7(nu447) +tbEx307*(mTruck), *mtx-2(gk444); miro-1(tm1966)+ tbEx307*(mTruck). n ≥25 animals for all genotypes, >250 long-lived actin for all genotypes. Kruskal Wallis ANOVA Dunn's test.

D) Quantitation of short-lived actin density/100µm/min in axons of HSN neuron of wild type and *ric-7(nu447)* mutants. n ≥25 animals. Mann-Whitney test.

E) Quantitation of trail density/100µm/min in axons of HSN neuron of wild type and *ric-7(nu447)* mutants. n ≥25 animals. Mann-Whitney test.

F) Quantitation of long-lived density/100µm/min in axons of HSN neuron of wild type and *ric-7(nu447)* mutants. n ≥25 animals. Mann-Whitney test.

G) Representative kymographs for EBP2::GFP from the major process of PLM neuron for wild type and *ric-7(nu447).* Scale bar: x axis= 5µm, y axis=15 sec.

H) Comparison between wild type and *ric-7(nu447)* for EBP2::GFP comets moving away from cell body (plus end out). n>20 animals.

I) Comparison between wild type and *ric-7(nu447)* EBP2::GFP comets moving cell body (minus end out) in the major process of PLM neuron. n>20 animals.

#### Supplementary Figure 3:

A) Representative GFP::UtCH kymograph for the proximal and distal region of PLM major process of *ric-7(nu447)* Purple and blue arrow head- representative trail in proximal and distal region respectively.

B) Representative GFP::UtCH kymograph for the proximal and distal region of PLM major process of wild type. Purple and blue arrow head- representative trail in proximal and distal region respectively.

C) Quantitation of short-lived actin density/100µm/min in proximal *ric-7(nu447)* and wild type. n>10 animals for both genotypes. Two-sample t-test with unequal variance, Welch correction.

D) Quantitation of short-lived actin density/100µm/min in distal region of *ric-7(nu447)* and wild type. n>10 animals for both genotypes. Two-sample t-test with unequal variance, Welch correction.

#### Supplementary Figure 4:

A) Schematic of CED-9-CED-3 pathway.

B) Representative kymograph for GFP::UtCH in PLM major process of wild type, Scale bar: x axis-5 µm, y axis-30 secs.

C) Representative kymograph for GFP::UtCH in PLM major process of *ric-7(nu447),* Scale bar: x axis-5 µm, y axis-30 secs.

D) Representative kymograph for GFP::UtCH in PLM major process of *ric-7(nu447)* *+tbEx306*(mTruck), Scale bar: x axis-5 µm, y axis-30 secs.

E) Representative kymograph for GFP::UtCH in PLM major process of *ced-3(n717),* Scale bar: x axis-5 µm, y axis-30 secs.

F) Representative kymograph for GFP::UtCH in PLM major process of *ced-3(n717); ric-7(nu447),* Scale bar: x axis-5 μm, y axis-30 secs.

G) Representative kymograph for GFP::UtCH in PLM major process of *ced-3(n717); ric-7(nu447) +tbEx306*(mTruck). Scale bar: x axis-5 μm, y axis-30 secs.

H) Quantitation of short-lived actin density/100μm/min in wild type, *sod-2(ok1030), sod-2(ok1030); ric-7(nu447), sod-2(ok1030); ric-7(nu447) +tbEx307*(mTruck). n=25 animals. Two-sample t-test for normally distributed data, Mann-Whitney test for non-normal data.

I) Quantitation of long-lived actin density/100μm/min in wild type, *sod-2(ok1030), sod-2(ok1030); ric-7(nu447), sod-2(ok1030); ric-7(nu447)*; + *tbEx307*(mTruck). n=25 animals. Two-sample t-test.

J) Quantitation of trail density/100μm/min for *ced-9(n2812), ced-3(n717), ced-9(n2812); ric-7(nu447), ced-3(n717); ric-7(nu447), ced-9(n2812); ric-7(nu447)*+ *tbEx306*(mTruck), *ced-3(n717); ric-7(nu447)*+ *tbEx306*(mTruck). n=20 animals. Two-sample t-test for normally distributed data, Mann-Whitney test for non-normal data.

K) Quantitation of short-lived actin density/100μm/min for *ced-9(n2812), ced-3(n717), ced-9(n2812); ric-7(nu447), ced-3(n717); ric-7(nu447), ced-9(n2812); ric-7(nu447)*+ *tbEx306*(mTruck), *ced-3(n717); ric-7(nu447)*+ *tbEx306*(mTruck). n=20 animals, Two-sample t-test for normally distributed data, Mann-Whitney test for non-normal data.

L) Quantitation of long-lived actin density/100μm/min for *ced-9(n2812), ced-3(n717), ced-9(n2812); ric-7(nu447), ced-3(n717); ric-7(nu447), ced-9(n2812); ric-7(nu447)*+ *tbEx306*(mTruck), *ced-3(n717); ric-7(nu447)*+ *tbEx306*(mTruck) n=20 animals. Two-sample t-test for normally distributed data, Mann-Whitney test for non-normal data.

M) Quantitation of mitochondrial density/100μm for *ced-9(n2812), ced-9(n2812); ric-7(nu447), ced-9(n2812); ric-7(nu447)*+ *tbEx306*(mTruck), *ric-7(nu447)*+ *tbEx306*(mTruck) n=20 animals. Two-sample t-test for normally distributed data, Mann-Whitney test for non-normal data.

N) Representative images of PH::roGFP_tsa2 in the AVA neuronal process of wild type at 488nm and 405 nm measuring cytosolic ROS.

O) Representative images of PH::roGFP_tsa2 in the AVA neuronal process of *ric-7(nu447)* at 488nm and 405 nm measuring cytosolic ROS.

P) Quantitation of ratio of PH::roGFP_tsa2 in 405/488 nm in the AVA neuronal processes of wild type and *ric-7(nu447).* Scale bar=5µm. n=10 animals. Two-sample t-test.

#### Supplementary Figure 5:

A) Distribution of proximal zone lengths of UNC-9::GFP in PLM neurons of wild type, *ric-7(nu447)*, *ric-7(nu447) +tbEx307*(mTruck), *ric-7(nu447) + twnEx337*[*mec-4*P::RHO-1(G14V)]).

B) Cluster size distribution of UNC-9::GFP in the PLM neurons of wild type, *ric-7(nu447)*, *ric-7(nu447) +tbEx307*(mTruck). n>30 puncta, N>10 animals. Kruskal Wallis ANOVA with Dunn's test.

C) Epifluorescence images of synaptogyrin at the cell body, region near the cell body and the chemical synaptic region of the PLM neuron in wild type. n>20 animals. Scale bar: x axis= 3µm.

D) Epifluorescence images of synaptogyrin at the cell body, region near the cell body and the chemical synaptic region of the PLM neuron in *ric-7(nu447)*. n>20 animals. Scale bar: x axis= 3µm.

E) Quantitation of trail density/100µm/min for wild type, *ric-7(nu447), ric-7(nu447)*+ *tbIs574*[$P_{TRN}$::RHO-1(G14V)] [$P_{TRN}$: *mec-4*p;TRN specific expression], wild type+ *tbIs574*[$P_{TRN}$::RHO-1(G14V)] [$P_{TRN}$: *mec-4*p;TRN specific expression]. n>25 animals. Kruskal Wallis ANOVA with Dunn's test.

F) Quantitation of mitochondrial density/100µm for wild type, *ric-7(nu447), ric-7(nu447)*+ *tbIs574*[$P_{TRN}$::RHO-1(G14V)] [$P_{TRN}$: *mec-4*p;TRN specific expression], wild type+ *tbIs574*[$P_{TRN}$::RHO-1(G14V)] [$P_{TRN}$: *mec-4*p;TRN specific expression]. n>25 animals. Kruskal Wallis ANOVA with Dunn's test.

G) Airy scan images of Spectrin in the PLM neurons of wild type, soluble GFP and *ric-7(nu447).* n>10 animals. Scale bar: x axis= 1µm.

H) Comparisons of periodicity in wild type and *ric-7(lf)*. one-way Kruskal Wallis ANOVA with Dunn's correction. n>10 animals.

### Supplementary movies

#### Movie 1: GFP::UtCH in PLM axon.

Time-lapse movie of GFP::UtCH in wild type PLM axon. Imaged sequentially at 1 frame per second (fps), playback at 60 fps. Genotype: *wyIs291* Cell body on left. Scale bar 10μm

#### Movie 2: GFP::UtCH in PLM axon of *ric-7(lf)*

Time-lapse movie of GFP::UtCH in PLM axon of *ric-7(lf)*. Imaged sequentially at 1 frame per second (fps), playback at 60 fps. Genotype: *ric-7(nu447)*; *wyIs291* Cell body on left. Scale bar 10μm

#### Movie 3: GFP::UtCH in PLM axon of *mtx-2(lf); miro-1(lf)*

Time-lapse movie of UtCH::GFP in PLM axon of *mtx-2(lf); miro-1(lf)*. Imaged sequentially at 1 frame per second (fps), playback at 60 fps. Genotype: *mtx-2(gk444); miro-1(tm1966)*. Cell body on left. Scale bar 10μm

#### Movie 4: GFP::UtCH in PLM axon of *ric-7(lf)* +mTruck

Time-lapse movie of GFP::UtCH in PLM axon of *ric-7(lf)* +mTruck. Imaged sequentially at 1 frame per second (fps), playback at 60 fps. Genotype: *ric-7(nu447); wyIs29; tbEx307*. Cell body on left. Scale bar 10μm

#### Movie 5: GFP::UtCH in PLM axon of *mtx-2(lf);miro-1(lf)*+ mTruck.

Time-lapse movie of GFP::UtCH in PLM axon of *mtx-2(lf);miro-1(lf)*+ mTruck. Imaged sequentially at 1 frame per second (fps), playback at 60 fps. Genotype: *mtx-2(gk444); miro-1(tm1966); wyIs291; tbEx307*. Cell body on left. Scale bar 10μm

#### Movie 6: GFP::UtCH in PLM axon of *ced-9(lf)*

Time-lapse movie of GFP::UtCH in PLM axon of *ced-9(lf)*. Imaged sequentially at 1 frame per second (fps), playback at 60 fps. Cell body on left. Genotype: *ced-9(n2812); wyIs291*. Scale bar 10μm

#### Movie 7: GFP::UtCH in PLM axon of *ced-3(lf)*

Time-lapse movie of UtCH::GFP in PLM axon of *ced-3(lf)*. Imaged sequentially at 1 frame per second (fps), playback at 60 fps. Cell body on left. Genotype: *ced-3(n717); wyIs291*Scale bar 10μm

#### Movie 8: GFP::UtCH in PLM axon of *ced-9(lf); ric-7(lf)*

Time-lapse movie of GFP::UtCH in PLM axon of *ced-9(lf); ric-7(lf)*. Imaged sequentially at 1 frame per second (fps), playback at 60 fps. Genotype: *ced-9(n2812); ric-7(nu447); wyIs291* Cell body on left. Scale bar 10μm

#### Movie 9: GFP::UtCH in PLM axon of *ced-3(lf); ric-7(lf)*

Time-lapse movie of GFP::UtCH in PLM axon of *ced-3(n717); ric-7(nu447); wyIs291*. Imaged sequentially at 1 frame per second (fps), playback at 60 fps. Cell body on left. Scale bar 10μm

#### Movie 10: GFP::UtCH in PLM axon of *ced-9(lf); ric-7(lf))* + mTruck

Time-lapse movie of GFP::UtCH in PLM axon of *ced-9(lf); ric-7(lf)* + mTruck. Imaged sequentially at 1 frame per second (fps), playback at 60 fps. Genotype: *ced-9(n2812); ric-7(nu447); wyIs291; tbEx306* Cell body on left. Scale bar 10μm

#### Movie 11: GFP::UtCH in PLM axon of *ced-3(lf); ric-7(lf)* + mTruck

Time-lapse movie of GFP::UtCH in PLM axon of *ced-3(lf); ric-7(lf)*+ mTruck. Imaged sequentially at 1 frame per second (fps), playback at 60 fps. Genotype: *ced-3(n717); ric-7(nu447); wyIs291; tbEx306* Cell body on left. Scale bar 10μm

#### Movie 12: GFP::UtCH in PLM axon of *sod-2(0)*

Time-lapse movie of GFP::UtCH in PLM axon of *sod-2(0)*. Imaged sequentially at 1 frame per second (fps), playback at 60 fps. Genotype: *sod-2(ok1030); wyIs291*. Cell body on left. Scale bar 10μm

#### Movie 13: UtCH::GFP in PLM axon of *sod-2(0); ric-7(lf)*

Time-lapse movie of GFP::UtCH in PLM axon of *sod-2(0); ric-7(lf)*. Imaged sequentially at 1frame per second (fps), playback at 60 fps. Genotype: *sod-2(ok1030); ric-7(nu447); wyIs291* Cell body on left. Scale bar 10μm

#### Movie 14: GFP::UtCH in PLM axon of *sod-2(0); ric-7(lf)* +mTruck

Time-lapse movie of GFP::UtCH in PLM axon of *sod-2(0); ric-7(lf)*. Imaged sequentially at 1frame per second (fps), playback at 60 fps. Genotype: *sod-2(ok1030); ric-7(nu447); wyIs291; tbEx307*. Cell body on left. Scale bar 10μm

#### Movie 15: EBP2::GFP in PLM neuron of wild type

Time-lapse movie of EBP2::GFP in PLM neuron of wild type. Imaged sequentially at 2 frames per seconds (fps), playback at 60 fps. Genotype: *juIs338*. Scale bar 10μm

#### Movie 16: EBP2::GFP in PLM neuron of *ric-7(lf)*

Time-lapse movie of EBP2::GFP in PLM neuron of *ric-7(lf)*. Imaged sequentially at 2 frames per seconds (fps), playback at 60 fps. Genotype: *juIs338; ric-7(nu447)*. Scale bar 10μm

### Supplementary tables

Table S1 Mean measurements from S1

| Properties of anterograde and retrograde trails | |
|---|---|
| | Mean ±SD |
| Anterograde trail velocity | 0.3µm/s ±0.13 |
| Retrograde trail velocity | 0.3µm/s ±0.13 |
| Anterograde trail length | 1.6µm ±0.94 |
| Retrograde trail length | 1.7µm ±0.88 |
| Anterograde trail density | 8.3 /100µm/min ±2.7 |
| Retrograde trail density | 8.5/100µm/min ±2.5 |

Table S2: Mean actin densities from figure 2 and S2

| Mean actin density/100µm/min ±SD | | |
|---|---|---|
| Genotype | Trail density/100mm/min | Short-lived density/100mm/min |
| Wild type (PLM) | 26/100 µm/min ±7.5 | 38.5/100 µm/min ±14.9 |
| *ric-7(lf) (*PLM*)* | 2.4/100µm/min ±3 SD | 0.3/100µm/min ±0.8 |
| *mtx-2(lf); miro-1(lf) (PLM)* | 2/100µm/min ±1.4 | 2.5/100 µm/min ±1.9 |
| Wild type (HSN) | 16.8/100µm/min ±6.2 | 44.8/100 µm/min ±15.3 |
| *ric-7(lf) (*HSN*)* | 2.5/100µm/min ±2.12 | 5.7/100 µm/min ±4.6 |

Table S3: Mean actin densities from figure 3 and S3

| Mean actin density/100µm/min ±SD | | | |
|---|---|---|---|
| Genotype | | Trail density/100µm/min | Short-lived density/100µm/min |
| Wild type | proximal | 15.1/100µm/min ±4.03 | 31.3/100µm/min ±10.55 |
| | distal | 21.5/100 µm/min ±5.5 | 38.5/100 µm/min ±8.0 |
| *ric-7(lf)* | proximal | 4.6/100µm/min ±3.8 | 7.4/100µm/min ±3.3 |
| | distal | 1.6/100 µm/min ±1.4 | 2.1/100 µm/min ±2.0 |

Table S4: Mean actin densities from figure 4 and S4

| Mean actin density/100µm/min ±SD | | |
|---|---|---|
| Genotype | Trail density/100µm/min | Short-lived density/100µm/min |
| Wild type | 19.2/100µm/min ±9.4 | 23.4/100µm/min ±9.8 |
| *ced-9(lf)* | 14.7/100µm/min ±5.9 | 24.3/100µm/min ±10.5 |
| *ced-9(lf); ric-7(lf)* | 6.9/100µm/min ±4.6 | 3.7/100µm/min ±3.2 |
| *ced-9(lf); ric-7(lf)* +mTruck | 23.0/100 µm/min ±10.38 | 27.0/100 µm/min ±12.7 |
| *ced-3(lf)* | 19.2/100µm/min ±7.4 | 32.4/100µm/min ±13.23 |
| *ced-3(lf); ric-7(lf)* | 1.8/100µm/min ±1.3 | 2.9/100µm/min ±3.5 |
| *ced-3(lf); ric-7(lf)* +mTruck | 19.0/100 µm/min ±7.0 | 39.5/100 µm/min ±15.2 |
| *sod-2(0)* | 25.9/100µm/min ±8.0 | 37.1/100µm/min ±8.6 |
| *sod-2(0); ric-7(lf)* (median) | 2.0/100 µm/min | 1.3/100 µm/min |
| *ric-7(lf)* (median) | 1.8/100 µm/min | 1.6/100 µm/min |
| *sod-2(0) ric-7(lf)* +mTruck | 9.2/100µm/min ±3.9 | 9.2/100µm/min ±3.9 |
| *ric-7(lf)* +mTruck | 23.8/100µm/min ±8.4 | 48.2/100µm/min ±15.3 |

Table S5 List of strains

| Sr No | Strain No. | Genotype | Reference/Source |
|---|---|---|---|
| 1 | N2 | Bristol wild type | (1) |
| 2 | TT2073 | *wyIs291* [*unc-86p*::GFP::UtCH; *odr-1p*::gfp] | (2) |
| 3 | TT2365 | *twnEx34* [*mec-7p*::mCherry::UtCH; *myo-2p*::GFP] | From Prof Chris Gabel |
| 4 | KP2048 | *ric-7(nu447)* | (3), CGC |
| 5 | VC1064 | *mtx-2(gk444)* | (4), CGC |
| 6 | TT1590 | *miro-1(tm1966)* | (5) |
| 7 | TT121 | *jsIs609* [*mec-7p*::MLS::GFP] | (6) |
| 8 | TT2540 | *tbEx307-* [*pmec-4::*UNC-116::tagRFP::TOMM7 (mTruck) (TTpl602) (20ng/µl*)* +*ttx-3p::*RFP (TTpl541) (50ng/µl), Isolate 1] | This study, plasmid gift from Josh Kaplan |
| 9 | TT2539 | *tbEx306 [pmec-4p::*UNC-116::tagRFP::TOMM7 (mTruck) (TTpl602*)* (20ng//µl*)* +*ttx-3p*::RFP (TTpl541) (50ng/µl, Isolate 2] | This study, plasmid gift from Josh Kaplan |
| 10 | TT2822 | *tbEx371* [*mec-4p*::UNC-116::tagRFP::TOMM7 (mTruck) (TTpl602) (3ng/µl) +*myo-2p*::GFP::H2B ( TTpl592) (50ng/µl)] | This study, plasmid gift from Josh Kaplan |
| 11 | TT1712 | *jsIs1073* | (7) |
| 12 | RB1072 | *sod-2(ok1030)* | (4), CGC |
| 13 | MT1522 | *ced-3(n717)* | CGC |
| 14 | MT4770 | *ced-9(n1950)* | CGC |
| 15 | TT3800 | *tbIs574*- integrated *tbEx529* [*mec-7p*::RHO-1(G14V)(TTpl772)(10ng/µl)+*myo2p*::mCherry (TTpl580)(25ng/µl) | This study, Plasmid was a gift from Chun-liang Pan |
| 16 | TT1428 | *twnEx337* [*mec-7p*::RHO-1 (G14V); *dpyp-30*::NLS::DsRed] | (8), gift from Chun-liang Pan |
| 17 | TT3827 | *ljEx437* [*mec-4p*::MEC-4::mCherry, *unc-122p*::GFP] | Gift from William Schafer, this study |
| 19 | TT3828 | *ljEx868* [*mec-4p*::UNC-7::GFP, *unc-122p*::mCherry] | Gift from William Schafer, this study |

| | | | |
|---|---|---|---|
| 20 | CZ18975 | *juIs338* [*mec-4p*::EBP-1::GFP +*ttx-3p*::RFP) | (9) |
| 21 | TT2884 | *tbIs388*-Integrated tbEx366 [*mec-4p*::SNG-1::GFP (TTpl696) (5ng/μl) + *myo-2p*::mCherry (TTpl580) (10ng/μl)] | (10) |
| 22 | TT3221 | *tbEx448[mec-4p::*UNC-9::GFP] | plasmid gift from Prof. Anindya Ghosh Roy |
| 23 | AQ3236 | *ljSi2* [*mec-7p*::GCaMP6m::SL2::TagRFP + *unc-119*(+)] | (11), CGC |
| 24 | MTS2000 | *spc-1(cas1047(spc-1::7xGFP11))(X); shyIs83 (mec-17p::GFP1-10; odr-1p::RFP* | (12) |
| 25 | MTS2501 | *spc-1(cas1047(spc-1::7xGFP11))(X); shyIs83 (mec-17p::GFP1-10; odr-1p::RFP)(I); n2657(ric-7(W192STOP(TGG>TGA))* | (12) |
| 26 | MTS2190 | *(shyEx534(mec-17p::3xGFP; odr-1p::GFP) (ex))* | (12) |
| 27 | FJH183 | *glr-1(ky176)*; *lin-15(n765ts)*; *csfEx61*[*rig-3p*::GLR-1::mCherry (pDM1556) (60ng/ul)+ *flp-18p*::PH::roGFP_tsa2(pRD14)(10ng/ul)+ LIN15+(pJM23)(25ng/ul)+ pBSKS(5ng/ul)] | Frederic Hoerndli |
| 28 | FJH719 | *glr-1(ky176)*; *ric-7(n2657*); *lin-15(n765ts)*; *csfEx61* | Frederic Hoerndli |

Table S6 List of plasmids

| Strain name | Details | Plasmid | Plasmid details | Reference |
|---|---|---|---|---|
| *tbEx307* | Generated by injecting TTpl602 (20ng/µl) with *ttx-3p::*RFP (TTpl541) (50ng/ml), as the co-injection marker | TTpl602 | UNC-116::tagRFP::TOMM7 including *unc-54* 3′-UTR was PCR amplified from PTT58 [unc129p::unc-116::tagRFP::tom7] using following primers, and cloned between Nhe-1 and Apa1 sites in TTpl503 [*mec4p*::LAMP-1::GFP] | Plasmid was a gift from Josh Kaplan, (13), this study. |
| *tbEx306* | Generated by injecting TTpl602 (20ng//µl) with *ttx-3p*::RFP (TTpl541) (50ng/µl) as the co-injection marker | TTpl602 | | |
| *tbEx371* | Generated by injecting TTpl602 (3ng/µl) with *myo-2p*::GFP::H2B ( TTpl592) (50ng/µl) as the co-injection marker | TTpl602 | | |
| *tbEx529* | Generated by injecting TTpl772 (10ng/µl) with *myo2p*::mCherry (TTpl580) (25ng/µl) as co-injection marker | TTpl772 | | Plasmid was a gift from Chun-liang Pan |

### References


1. S. Brenner, The genetics of Caenorhabditis elegans. *Genetics* **77**, 71-94 (1974).
2. P. H. Chia, B. Chen, P. Li, M. K. Rosen, K. Shen, Local F-actin network links synapse formation and axon branching. *Cell* **156**, 208-220 (2014).
3. Y. Hao, Z. Hu, D. Sieburth, J. M. Kaplan, RIC-7 promotes neuropeptide secretion. *PLoS Genet* **8**, e1002464 (2012).
4. Anonymous, large-scale screening for targeted knockouts in the Caenorhabditis elegans genome. *G3 (Bethesda)* **2**, 1415-1425 (2012).
5. G. R. Sure *et al.*, UNC-16/JIP3 and UNC-76/FEZ1 limit the density of mitochondria in C. elegans neurons by maintaining the balance of anterograde and retrograde mitochondrial transport. *Sci Rep* **8**, 8938 (2018).
6. C. Fatouros *et al.*, Inhibition of tau aggregation in a novel Caenorhabditis elegans model of tauopathy mitigates proteotoxicity. *Hum Mol Genet* **21**, 3587-3603 (2012).
7. Q. Zheng *et al.*, The vesicle protein SAM-4 regulates the processivity of synaptic vesicle transport. *PLoS Genet* **10**, e1004644 (2014).
8. C. H. Chen, C. W. He, C. P. Liao, C. L. Pan, A Wnt-planar polarity pathway instructs neurite branching by restricting F-actin assembly through endosomal signaling. *PLoS Genet* **13**, e1006720 (2017).
9. M. Chuang *et al.*, The microtubule minus-end-binding protein patronin/PTRN-1 is required for axon regeneration in C. elegans. *Cell Rep* **9**, 874-883 (2014).
10. S. S. P. Nadiminti *et al.*, Active zone protein SYD-2/Liprin-α acts downstream of LRK-1/LRRK2 to regulate polarized trafficking of synaptic vesicle precursors through clathrin adaptor protein complexes. *bioRxiv* (2023).
11. Y. Cho *et al.*, Automated and controlled mechanical stimulation and functional imaging in vivo in C. elegans. *Lab Chip* **17**, 2609-2618 (2017).
12. O. Glomb *et al.*, A kinesin-1 adaptor complex controls bimodal slow axonal transport of spectrin in Caenorhabditis elegans. *Dev Cell* **58**, 1847-1863.e1812 (2023).
13. T. Zhao, Y. Hao, J. M. Kaplan, Axonal Mitochondria Modulate Neuropeptide Secretion Through the Hypoxic Stress Response in Caenorhabditis elegans. *Genetics* **210**, 275-285 (2018).
